# Generalizable approaches for genomic prediction of metabolites in plants

**DOI:** 10.1101/2021.11.24.469870

**Authors:** Lauren J. Brzozowski, Malachy T. Campbell, Haixiao Hu, Melanie Caffe, Lucía Gutiérrez, Kevin P. Smith, Mark E. Sorrells, Michael A. Gore, Jean-Luc Jannink

## Abstract

Plant metabolites are important for plant breeders to improve nutrition and agronomic performance, yet integrating selection for metabolomic traits is limited by phenotyping expense and limited genetic characterization, especially of uncommon metabolites. As such, developing biologically-based and generalizable genomic selection methods for metabolites that are transferable across plant populations would benefit plant breeding programs. We tested genomic prediction accuracy for more than 600 metabolites measured by GC-MS and LC-MS in oat (*Avena sativa* L.) seed. Using a discovery germplasm panel, we conducted metabolite GWAS (mGWAS) and selected loci to use in multi-kernel models that encompassed metabolome-wide mGWAS results, or mGWAS from specific metabolite structures or biosynthetic pathways. Metabolite kernels developed from LC-MS metabolites in the discovery panel improved prediction accuracy of LC-MS metabolite traits in the validation panel, consisting of more advanced breeding lines. No approach, however, improved prediction accuracy for GC-MS metabolites. We tested if similar metabolites had consistent model ranks and found that, while different metrics of ‘similarity’ had different results, using annotation-free methods to group metabolites led to consistent within-group model rankings. Overall, testing biological rationales for developing kernels for genomic prediction across populations, contributes to developing frameworks for plant breeding for metabolite traits.

## INTRODUCTION

Plant metabolites contribute to human health, food flavor, and plant resistance to stresses, and thus are important traits for plant breeders (Kumar et al., 2017; Zhu et al., 2019). While selection for some metabolites is possible through correlated traits, like color, many metabolites are phenotyped through metabolomics approaches like chromatography and mass spectrometry (Fernie & Tohge, 2017). Some key challenges in plant breeding for metabolites are the diversity of plant metabolites, with hundreds of thousands predicted (Afendi et al., 2012), a generally limited knowledge of the genetic architecture of metabolite traits (Soltis & Kliebenstein, 2015), and expense in generating metabolomics data. As our capacity to measure and identify plant metabolites grows (Fernie & Tohge, 2017), developing biologically-based and generalizable selection methods that are transferable across plant populations would benefit plant breeding programs.

Most knowledge of the genetic bases of metabolite variation in crops comes from models like tomato, maize, and rice, and nutritional metabolites, such as vitamin precursors (Luo, 2015; Fernie & Tohge, 2017; Wager & Li, 2018). While this work encompasses biochemical pathways that are largely conserved, there is also a growing body of work on specialized metabolites, metabolites that contribute to ecological interactions and are generally restricted to few lineages, for instance, alkaloid production in tomato (Zhu et al., 2018) and benzoxazinoid production in maize (Zhou et al., 2019). Together, these studies have shaped our understanding of the genetic architecture of plant metabolite traits: while some specialized metabolites have oligogenic genetic architecture (Diepenbrock et al., 2017, 2021), many loci contributing to metabolite variation have small effects, and there are multiple examples of balancing selection for metabolites (Soltis & Kliebenstein, 2015). Given the typically complex genetic architecture and small-effect loci that underpin metabolite traits, techniques like genomic prediction and selection would be particularly useful methods to implement in plant breeding programs (Heffner et al., 2009; Heslot et al., 2015).

Genomic prediction and selection studies have shown that metabolomic traits are viable candidates for genomic selection. For instance, genomic selection on color (a proxy for provitamin A) in winter squash (*Cucurbita moschata*) fruit, led to significant population improvement over four cycles of selection (Hernandez et al., 2020). In addition, average genomic prediction accuracy for measured vitamin metabolites was 0.43 for provitamin A in maize kernels (Owens et al., 2014), and 0.49 for vitamin E in fresh sweet corn kernels (Baseggio et al., 2019). Recently, others have also tested strategies for incorporating multiomic information in prediction of metabolites. For instance, computing relationship matrices from metabolomics data (Campbell et al., 2021a) or metabolomics and transcriptomics data (Hu et al., 2021) led to high average prediction accuracies (r > 0.4) for fatty acid traits in oat (*Avena sativa*) seed. These studies have demonstrated that genomic prediction is effective for a few to tens of biochemically similar metabolites traits. Expanding to consider more metabolites would allow for an understanding of the generalizability of the results. Further, as with much work involving multiomic datasets, connecting genomic prediction results to biological mechanisms is a challenge.

One approach to elucidate and incorporate biological bases into genomic prediction has been through tests of genomic partitioning where, if the partitioned SNPs are enriched for causal variants, prediction accuracy could be improved (Sarup et al., 2016). Recent work in genomic prediction of 65 free amino acid metabolite traits in *Arabidopsis* seeds partitioned genomic SNPs using annotations from 20 biochemical pathways, and found that inclusion of pathway SNPs as a kernel in a multikernel BLUP model improved prediction ability (Turner-Hissong et al., 2020). In other examples, genomic prediction with pathway SNPs alone was equivalent to genome-wide prediction for provitamin A compounds (carotenoids) in maize kernels (Owens et al., 2014), but biosynthetic pathway SNPs performed worse than genome-wide SNPs for prediction of vitamin E (tocochromanols) in fresh sweet corn kernels (Baseggio et al., 2019). These differences could be due to the degree to which markers were in LD with causal variants (Baseggio et al., 2019) or may point to causal variation being attributable to regulation (local or distal), or factors like metabolite transport (Soltis & Kliebenstein, 2015). Finally, while integrating prior information about biochemical pathways has promising but mixed success, its application remains limited to organisms with well annotated genomic, transcriptomic and metabolomic resources.

Strategies to conduct genomic partitioning without incorporating prior biosynthesis information have also been tested. In oat (*Avena sativa* L.), a hexaploid with a recently available whole genome sequence, (Campbell et al., 2021b) leveraged untargeted metabolomics data with over 1600 metabolites to conduct factor analysis to uncover genomic regions that influence metabolite composition. Using a multi-kernel approach, incorporating a kernel using GWAS results of factors improved prediction accuracy of lipid and protein traits across populations (Campbell et al., 2021b). In this analysis, factors were most commonly enriched for lipids which perhaps contributed to increased prediction accuracy of fatty acids (a type of lipid), but it would be intriguing to understand if this result is generalizable across more types of metabolites that were less represented in the factor data set.

We sought to expand upon the work of (Campbell et al., 2021b) to test prediction models for the entire oat seed metabolome and develop generalized genomic prediction method frameworks. Oat seeds contain multiple healthful metabolites such as unsaturated fatty acids, beta-glucans, fiber as well as antioxidants (Stewart & McDougall, 2014), and fatty acid traits have been a target of GWAS (Carlson et al., 2019) and genomic prediction (Campbell et al., 2021b; a; Hu et al., 2021). Using this well-studied germplasm, we examined more than 600 metabolites in oat seed measured by GC-MS and LC-MS and tested genomic prediction accuracy using two-kernel models. Our objectives were to characterize the measured metabolome by metabolite GWAS (mGWAS), leverage mGWAS results to select loci for two-kernel genomic prediction models to test hypotheses about the most informative, biologically-based genome partitioning methods of metabolomics data, and to evaluate prediction accuracy of these models in a separate germplasm panel. To this end, we conducted mGWAS in a discovery panel and generated kernels from significant mGWAS SNPs for any metabolite, or of metabolites identified by structure as lipids or belonging to specific biosynthetic pathways thereof (terpenoid biosynthesis pathways). Genomic prediction accuracy was evaluated in a validation germplasm panel using K-fold cross validation. We hypothesized that kernels encompassing metabolome-wide information would increase prediction accuracy for many metabolites, while kernels for specific metabolite types or pathways would result in the highest prediction accuracy of their own metabolites. We also hypothesized that similar metabolites would have similar genomic prediction results (in terms of model rank), and defined metabolomic ‘similarity’ in three ways: high-confidence annotations, structural annotations, or by an annotation-free method. Broadly, as plant breeders target larger numbers and more diverse (less well known) metabolites, developing frameworks for structuring genomic prediction models is important. This work tests different biological rationales for incorporating information into genomic prediction, and transferability across populations.

## MATERIALS and METHODS

### Oat metabolome discovery phenotypes

Whole metabolome phenotypes were measured from mature seeds using untargeted LC-MS and GC-MS in a diverse oat germplasm panel of 375 inbred lines. These phenotypes have been previously described (Brzozowski et al., 2021; Campbell et al., 2021b; a; Hu et al., 2021). For each metabolite phenotype, measured as relative signal intensity, deregressed best linear unbiased predictors (drBLUPs) could be calculated for 1067 of the LC-MS and 601 of the GC-MS signals as in (Campbell et al., 2021b).

We characterized the metabolites by information provided by the Proteomics and Metabolomics Facility at Colorado State University (Fort Collins, CO, USA) (Table 1). The metabolites were annotated by comparison to an in-house spectral library RAMSearch (Broeckling et al., 2016) and MSFinder (Tsugawa et al., 2016), and details of measurement and annotation of this dataset are provided in (Brzozowski et al., 2021; Campbell et al., 2021b; a; Hu et al., 2021). To further characterize the metabolites, we examined the continuous variables of retention time (a measure of polarity, where, using a reverse phase column, a lower retention time indicates greater polarity), molecular mass, and genomic heritability (de los Campos et al., 2015). We also used the provided categorical variables of instrument type (LC, GC), and multiple levels of metabolite type as identified by ClassyFire (Djoumbou Feunang et al., 2016) by superclass, class and subclass. ClassyFire was run in the ClassyFire Batch Compound Classification web server (https://cfb.fiehnlab.ucdavis.edu/) on July 1, 2021.

**Table 1.** Metabolite classification of the discovery panel for categorical variables of ClassyFire superclass and class, and numeric metrics of retention time and molecular mass. The distribution of metabolite retention time and molecular mass are given in **Figure S1**.

| Classification | LC | GC | Total |
| --- | --- | --- | --- |
| **ClassyFire Classification (count)** |  |  |  |
| ***Lipids and lipid-like molecules (“Lipids”)*** | **527** | **6** | **533** |
| ...Glycerophospholipids | 123 | 1 | 124 |
| ...Glycerolipids | 87 | 0 | 87 |
| ...Fatty Acyls | 99 | 5 | 104 |
| ...Steroids and steroid derivatives | 83 | 0 | 83 |
| ...Prenol lipids | 102 | 0 | 102 |
| ***Organoheterocyclic compounds (“OgHc”)*** | **57** | **9** | **66** |
| ***Phenylpropanoids and polyketides (“Phe”)*** | **67** | **2** | **69** |
| ...Cinnamic acids and derivatives | 13 | 1 | 14 |
| ...Coumarins | 11 | 0 | 11 |
| ***Organic acids and derivatives (“OgAc”)*** | **70** | **26** | **96** |
| ...Carboxylic acids and derivatives | 41 | 22 | 63 |
| ***Organic oxygen compounds (“OgOx”)*** | **68** | **20** | **88** |
| ***Other<sup>1</sup>*** | **74** | **6** | **80** |
| ***Not classified (“None”)*** | **204** | **532** | **736** |
| **Numeric metrics (mean)** |  |  |  |
| Retention time (s) | 515.4 | 731.6 | 593.3 |
| Molecular mass (g/mol) | 669.8 | NA | NA |

<sup>1</sup>The ‘Other’ classification includes the nucleosides, nucleotides and analogues, and organic nitrogen compounds superclasses for all metabolites, and the alkaloids and derivatives, hydrocarbons, lignans, neolignans and related compounds, organic polymers and organosulfur compounds, and benzenoids superclasses for LC metabolites and homogenous non-metal for GC metabolites

### Genomic analysis of discovery panel metabolome

All analyses were conducted in the R programming environment (R Core Team, 2016). We obtained genotyping-by-sequencing (GBS) data from T3/Oat (https://oat.triticeaetoolbox.org/) for 342 individuals in a diverse panel of oat genotypes as described in (Campbell et al., 2021b). The GBS data was filtered (less than 40% missingness, minor allele frequency greater than 0.02) and imputed with glmnet (Friedman et al., 2010). Of these 73,527 markers, the 54,284 that could be anchored to the genome (PepsiCO OT3098v1; https://wheat.pw.usda.gov/GG3/graingenes_downloads/oat-ot3098-pepsico) were used. A principal component (PC) analysis was conducted using the centered and scaled matrix of allele dosages with the function ‘prcomp’, and percent variance explained by each PC was found using the ‘fviz_eig’ function. By examination of the scree plot, the first five PCs (accounting for 21.8% of the variance) were chosen for use in analysis. A kinship matrix was calculated using the ‘A.mat’ function, and genomic heritability was calculated using variance components extracted from the ‘kin.blup’ function, both in the R package rrBLUP (Endelman, 2011).

### Genome wide association study in discovery panel

A single-trait genome-wide association study was conducted for all metabolites (mGWAS) in the statgenGWAS package (Rossum & Kruijer, 2020) using the kinship matrix and using five PCs as covariates. A false discovery rate correction was used on *p*-values for each metabolite, and a result was considered significant if *p_FDR_* < 0.05.

### Defining metabolite kernels from discovery panel

We defined sets of SNPs that may broadly shape the measured seed metabolome (“general” kernels), and those that are more specific to lipids (“lipid” kernels) (Table 2). For general kernels, we selected SNPs that were significant mGWAS results for: (1) three or more LC or GC metabolites (“Any3”), (2) at least one LC metabolite and at least one GC metabolite (“LCGC2”), (3) four or more LC metabolites (“LC4”), or (4) two or more GC metabolites (“GC2”). The different criteria used to construct LC4 and GC2 were chosen to compare a similar proportion of metabolites per instrument (0.37% and 0.33%, respectively). To determine if these kernels represented more SNPs than expected by chance, we used a Poisson model to determine the probability of observing the same significant SNP for multiple metabolites to the rate of SNP inclusion in a kernel using the ‘ppois’ function in R.

**Table 2.** Description and prediction of performance of metabolite kernels. The groups “MEP” and “MVA” refer to the Methylerythritol Phosphate pathway and Mevalonate Acid pathway branches of terpenoid biosynthesis, respectively.

| *Type* | *Group* | *Description* | *Rationale* | *Predictions* |
| --- | --- | --- | --- | --- |
| General metabolome | Any3 | GWAS results shared by any three or more metabolites (LC or GC) | Metabolites from either instrument, extraction method contribute equally to capturing broader metabolome variation | Since both instruments are included, will perform best for a broad range of metabolites |
|  | LCGC2 | GWAS results shared by at least one LC and at least one GC metabolite | Including metabolites from both instruments, extraction methods, is necessary to capture metabolome variation |  |
|  | LC4 | GWAS results shared by four or more LC metabolites | Metabolites from a single instrument, extraction method, but not restricted to a specific class | Will perform better for metabolites from respective instruments, but will still perform well for a broad range of metabolites |
|  | GC2 | GWAS results shared by two or more GC metabolites | Metabolites from a single instrument, extraction method, but not restricted to a specific class |  |
| Lipids | Lipid | GWAS results shared by two or more LC lipids | Metabolites from a single instrument, extraction, restricted to lipids | Will perform well for lipids, but increased specificity (MVA, MEP) will reduce performance for metabolites overall |
|  | MVA, and MEP | GWAS results of any LC MVA- or MEP-derived terpenoids | Metabolites from a single instrument, extraction method, restricted to specific biosynthetic pathways of terpenoids |  |

We defined lipid kernels by significant mGWAS results of LC-MS lipids based on a hierarchy of pathway specificity. First, we defined a kernel of significant mGWAS results shared by two or more metabolites classified as ‘Lipids and lipid-like molecules’ superclass (“Lipid”). We also created two terpenoid biosynthesis pathway kernels of significant mGWAS results from metabolites classified as (1) the subset of terpenoids predominantly produced by the Mevalonate Acid pathway (“MVA”; subclasses of ‘Triterpenoids’ and ‘Sesquiterpenoids’), and (2) the subset of terpenoids predominantly produced by the Methylerythritol Phosphate pathway (“MEP”; subclasses of ‘Diterpenoids’ and ‘Tetraterpenoids’). Again, criteria for including SNPs were modified by kernel to create kernels of similar size.

We visualized genome location by plotting the number of significant mGWAS results in 10Mb bins. For all further analyses we added all other SNPs in strong linkage disequilibrium LD (*r^2^*>0.5) to each set of SNPs. We used the most recent transcriptome annotations (Hu et al., 2020) and noted SNPs that were within, or up to 2.5kb upstream of genes.

### Descriptive analyses of metabolite kernels

We examined if metabolite characteristics were explanatory for the GWAS results identified. First, we tested if there was a relationship between metabolite heritability and retention time, molecular mass, or metabolite superclass. For retention time and molecular mass, we used a linear model with the ‘lm’ function with heritability as the response variable, and tested effect significance by ANOVA. We also calculated mean heritability for metabolites by ClassyFire superclass.

We tested if focal superclass categories were enriched or depleted in each of the kernels using the ‘phyper’ function in R. We also calculated the mean Euclidean distance between metabolites in the kernels, using a matrix with metabolites in rows and oat lines in columns and the cells containing their scaled and centered drBLUPs with the ‘dist’ function with the ‘euclidean’ method in R. To compare distance between metabolites contributing to the kernel to metabolites not contributing to the kernel, we used the Mann-Whitney *U* test implemented with the ‘wilcox.test’ function in R.

### Oat metabolome validation phenotypes

We used a validation germplasm panel to test the transferability of kernels between populations. This population is described by (Brzozowski et al., 2021). Briefly, a panel of 235 inbred lines was evaluated in three Midwest production environments (Minnesota, “MN”; South Dakota, “SD” and Wisconsin, “WI”). For this analysis, we removed lines that overlapped with our discovery (diverse) panel, leaving 212 lines in MN and SD and 208 lines in WI. The relationship between the discovery and validation panels are described in (Hu et al., 2021), named as ‘discovery’ and ‘elite’ panels, respectively.

Deregressed BLUP (drBLUP) were calculated as in (Campbell et al., 2021b) where data was cube-root transformed, and there were 397 LC and 243 GC metabolites (640 total) for which drBLUPs could be calculated. Metabolite heritability and percent variation described by kernels were calculated as above. Spearman’s rank correlation of metabolite heritability across environments was evaluated with the ‘cor.test’ function in R. In addition to examining the metabolome as a whole, we also evaluated outcomes for the specialized metabolites, avenanthramides, avenacins and avenacosides as described in (Brzozowski et al., 2021). Metabolite drBLUPs and annotations are provided as **Supporting Data.**

### Genomic prediction in the validation panel

We conducted genomic prediction for metabolites (*n*=640) and genotypes (*n*=189) measured in the validation panel separately in all environments. We then fit a two-kernel GBLUP model using the selected SNPs to construct Gaussian kernels as described in (de Los Campos, 2018) and (Cuevas et al., 2020) in the R package ‘BGLR’ (Pérez & de los Campos, 2014) with 20000 iterations and a burn in of 5000. We conducted five-fold cross validation with 50 replicates, where folds were consistent between metabolites and environments, and report the correlation (*r*) between predicted and observed values.

### Evaluation of genomic prediction results in the validation panel

We evaluated if the two kernel metabolite models had significantly higher or lower prediction accuracies than GBLUP. First, we used paired one-sided Wilcoxon rank-sum tests using the mean prediction accuracy per metabolite and model. We also tested if mean prediction accuracy varied between environments using a Kruskal-Wallis test. Finally, we conducted paired tests between the two-kernel metabolite models and GBLUP by models and metabolites using accuracy of each of fifty replicates to understand which were significantly different from GBLUP. In both cases, we report significant results as *p_BONF_*< 0.05.

We partitioned genetic variation from the two kernels (metabolite kernel, rest-of-genome kernel) to assess the percent variation that was explained by the metabolite kernels. We compared the metabolite kernels described above to kernels constructed from random draws of loci with significant mGWAS results that were not included in metabolite kernels (*n*=4238 SNPs). We had 10 random draws of 20, 50, 100, 500, 900 and 1800 SNPs, and added SNPs in LD as above to span the size range of kernels (**Table S3**). The genetic variation explained by these null kernels relative to metabolite kernels was evaluated as well as the impact of kernel size on genetic variation explained.

To examine differences between environments, we created matrices of prediction accuracies with models in rows and each metabolite in columns by environment. We then calculated the distance between models (by metabolites of each instrument) and performed hierarchical clustering within an environment and compared groupings of models.

Finally, we tested if similar metabolites have similar model ranks, measured by Spearman’s rank correlation. We defined ‘similar’ in three ways. First, we examined results for seven specialized metabolites important for human health, or plant resistance to disease for which we have high-confidence annotations: the avenanthramides, avenacins and avenacosides (Brzozowski et al., 2021). Second, we used finer scale structural descriptions (‘Class’ description) of metabolites of the ‘Lipid and Lipid-like compounds’ ClassyFire Superclass (*n*=91). Third, we attempted an annotation-free method where we computed the mean Euclidean distance between metabolites in the kernels with metabolites in rows and oat lines in columns and the cells containing their scaled and centered drBLUPs with the ‘dist’ function with the ‘euclidean’ method in R. We then performed hierarchical clustering to define 10 groups of metabolites for each of the environments using the ‘hclust’ function both in R.

## RESULTS

### Oat seed metabolome of the discovery panel

Using untargeted metabolomics, we detected 1067 LC-MS and 601 GC-MS metabolites for which deregressed BLUPs could be calculated, and characterized the metabolites by chemical properties as well as retention time and molecular mass. The LC-MS metabolites had greater genomic heritability (mean, *h^2^*=0.23) than GC-MS metabolites (mean, *h^2^*=0.13) (Figure 1a). For both LC-MS and GC-MS metabolites, we found that heritability was greater at lower retention times (greater polarity) and for larger molecular masses, even when the lowest heritability compounds were excluded (**Figure S1**). The LC-MS metabolites were more densely annotated than the GC-MS metabolites, and lipids were the most common classification (49%) of LC-MS metabolites (Table 1). While we did not observe any relationship between heritability and metabolite structural characteristics, annotated GC-MS metabolites had higher heritability than unannotated metabolites (**Table S1**).

**Figure 1.**
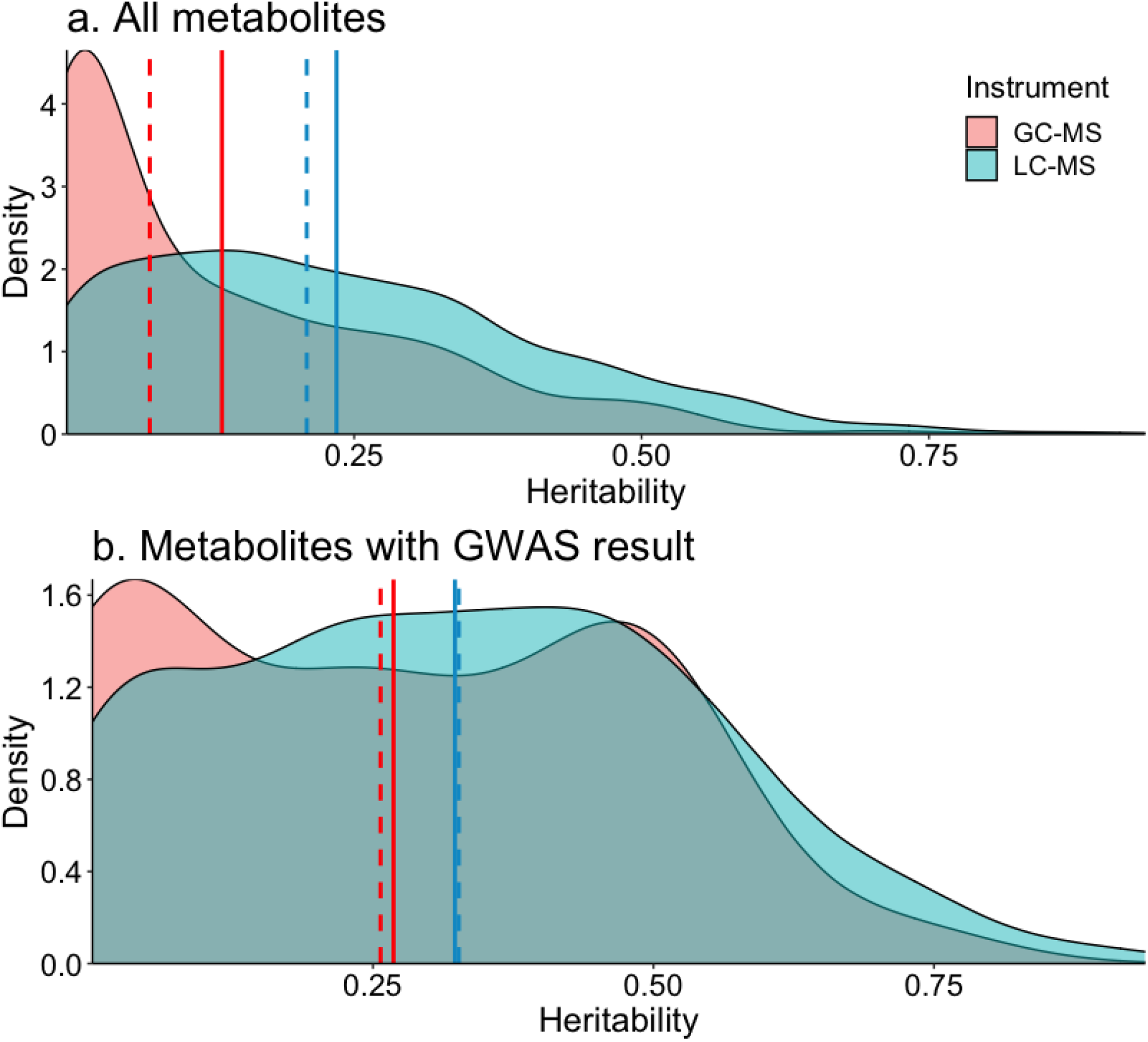
Genomic heritability (a) all metabolites (*n*=1668) and (b) metabolites with a significant GWAS (*n*=368) result from the discovery panel. The instrument class (LC-MS, or GC-MS) is denoted by color (blue, red, respectively). The solid line indicates the mean and dashed line indicates the median genomic heritability by instrument class.

A metabolite genome-wide association study mGWAS was conducted for all metabolites, and 368 metabolites had at least one significant SNP (*p_FDR_* < 0.05) and 8415 unique SNPs (15.5% of total SNPs) were implicated. Of these, there were 282 LC-MS (5728 unique SNPs, 10.6% of total SNPs), and 86 GC-MS (3544 unique SNPs, 6.5% of total SNPs) metabolites with a significant association. The metabolites with significant associations tended to have higher heritability than those without for both LC-MS and GC-MS metabolites (Figure 1b**; Figure S2**).

### Defining kernels for whole genome regression

Using the mGWAS results, we defined kernels to capture loci that broadly shape the metabolome (“general”), and loci specific to metabolite structures or pathways. We hypothesized that the general kernels would broadly improve metabolite prediction, while kernels customized to specific lipids would improve prediction of their respective metabolites (Table 2).

The kernels included 493-1800 and 109-917 significant mGWAS SNPs from 60-274 and 9-78 metabolites for the general and specific kernels, respectively (**Table S2**), with some metabolites and SNPs contributing to multiple kernels (**Figure S3)**. Correlations between kernel off-diagonal elements ranged from *r*=0.12 - 0.83, and the two kernels relying on mGWAS from GC-MS (‘LCGC2’ and ‘GC2’) were the most distinct from other kernels (**Figure S4**).

In evaluating if kernels were enriched for mGWAS loci from particular metabolites, we found that LC-MS metabolites contributing to metabolite kernels were significantly depleted for lipids (Figure 2). GC-MS metabolites were more sparsely annotated than LC-MS compounds, but metabolites with mGWAS results were enriched for annotated compounds (Figure 2). We also evaluated the pairwise Euclidean distance between metabolites to test in an annotation-free way if more similar metabolites had similar mGWAS results. The GC-MS metabolites contributing to kernels had significantly reduced distance between metabolites compared to all GC-MS metabolites, but there was no reduced distance of LC-MS metabolites contributing to kernels (**Figure S5**).

**Figure 2.**
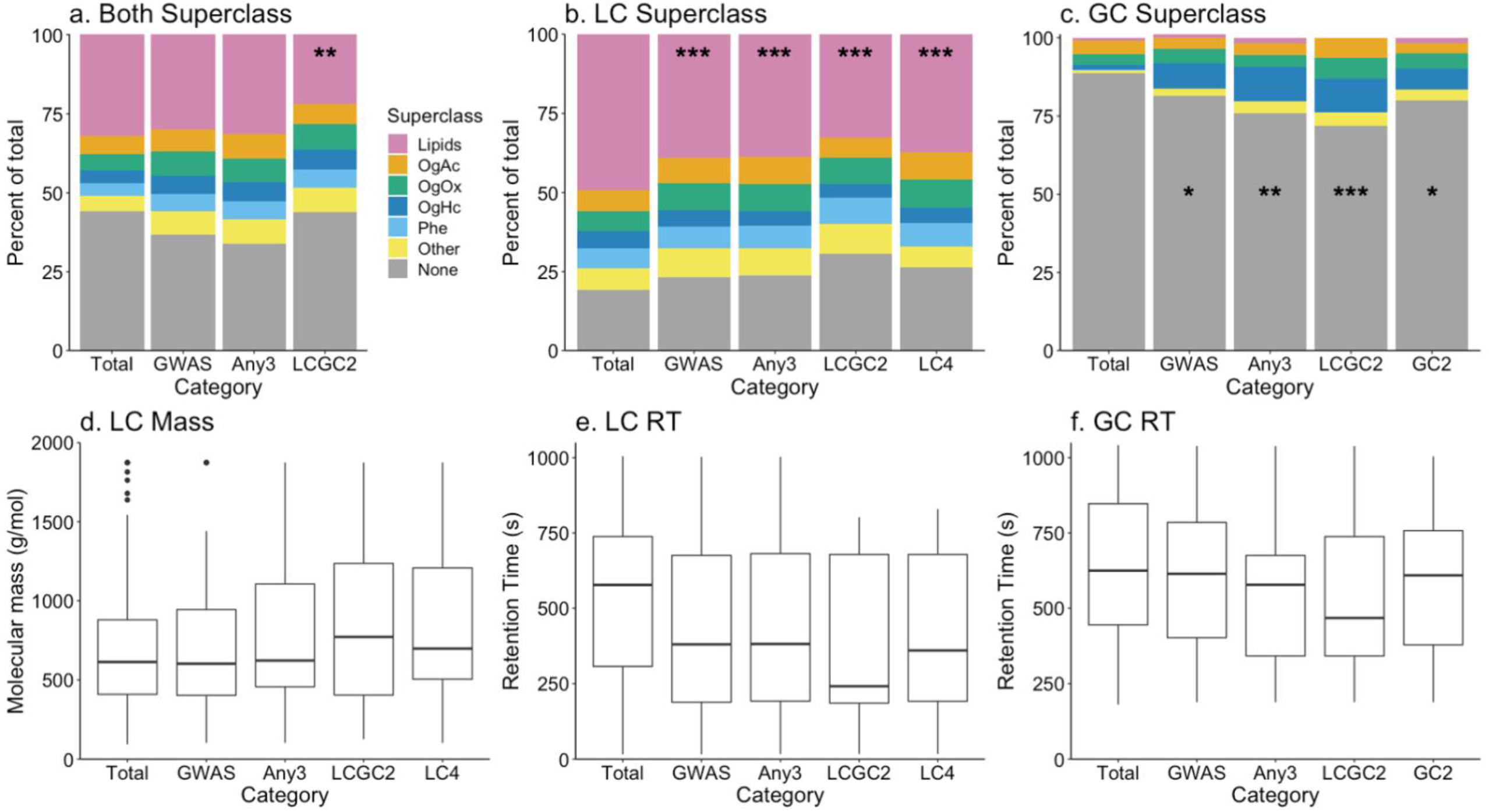
Distribution of metabolites by ClassyFire Superclass by general metabolite kernel in the discovery panel for (a) both LC-MS and GC-MS metabolites, (b) LC-MS metabolites only and (c) GC-MS metabolites only. Distributions of (d) LC-MS molecular mass, and (e) LC-MS and (f) GC-MS retention time (“RT”) are shown by kernel. Significance indicators identify instances of depletion where * *p*<0.05, and ** *p<*0.01 and *** *p*< 0.001. Abbreviations of metabolite superclass are given in Table 1.

We compared the rate of SNPs meeting criteria for inclusion in a kernel (e.g., significant mGWAS result shared by three metabolites) to the empirical rate of mGWAS results in this oat population. Compared to a random draw from a Poisson distribution, there were more SNPs meeting criteria than expected (‘Any3’, λ=0.16, *p*= 5.5e-04; ‘LCGC2’, λ=0.16, *p*= 1.2e-04; ‘LC4’, λ=0.11, *p*= 4.8e-06; ‘GC2’, λ=0.07, *p*=0.002). The SNPs for the general kernels were identified on most chromosomes but clustered within chromosomes (**Figure S6**). The lipid-related kernels had the most SNPs on chromosome 5A and 5C (**Figure S7**). Finally, kernels had a range of gene density, with a maximum 11% of SNPs in the ‘MVA’ kernel being in a gene and a minimum of 6.7% in ‘LCGC2’ (Table 3).

**Table 3.** Number of genes associated with each metabolite kernel. Kernel size is given in **Table S3**. The total genes implicated (‘total genes’), the number of genes per SNP in the kernel (‘genes per SNP’), and percent of SNPs in kernel in a gene (‘Percent SNPs with a gene’) are shown.

| Kernel | Total genes | Genes per SNP | Percent SNPs within gene |
| --- | --- | --- | --- |
| MVA | 127 | 0.127 | 11.01 |
| LC4 | 261 | 0.101 | 8.63 |
| Lipid | 225 | 0.092 | 8.07 |
| Any3 | 455 | 0.086 | 7.72 |
| MEP | 53 | 0.085 | 6.77 |
| GC2 | 150 | 0.072 | 6.80 |
| LCGC2 | 183 | 0.071 | 6.67 |

### Oat seed metabolome of the validation panel

We tested if kernels developed in the discovery panel improved prediction accuracy for metabolites in a validation panel evaluated in three environments (Minnesota, “MN”; South Dakota, “SD” and Wisconsin, “WI”) that had 397 LC-MS and 243 GC-MS metabolites. Although the measurements do not allow for direct comparison of all individual metabolites to those in the discovery panel (due to currently no robust method to map all untargeted metabolites from one panel to another and quantify them accurately, Hu *et al.* 2021), the metabolite classification parameters were consistent across the two panels. Like the discovery panel, LC-MS metabolites had greater mean heritability (*h^2^*: MN=0.30, SD=0.17, WI=0.17) than GC-MS metabolites (*h^2^*: MN=0.10, SD=0.09, WI=0.14) and heritability was positively correlated across environments (**Table S4**). Metabolite classifications were available for the LC-MS metabolites only, and lipids were the most common annotation (23%), but there were no trends in heritability by metabolite type (**Table S5**). Finally, except for LC-MS metabolites in MN, there were significant negative relationships between heritability and retention time (**Figure S8**, **Table S6**).

### Genomic prediction in the validation panel

Mean prediction accuracy of two-kernel (metabolite kernel and rest-of-genome kernel) genomic prediction models from five-fold cross validation ranged from 0.24-0.34 for LC-MS and 0.13-0.17 for GC-MS metabolites, where prediction accuracy was highest for LC-MS metabolites in MN and lowest for GC-MS metabolites in MN and SD (Table 4). The ‘LC4’ kernel improved and the ‘GC2’ kernel reduced prediction accuracy of LC-MS metabolites over GBLUP in all three environments (Figure 3a). The ‘Any3’ kernel also improved prediction accuracy of LC-MS metabolites over GBLUP in two environments, as did the ‘MVA’ kernel, contrary to our expectation that the ‘MVA’ kernel specificity would not result in improved prediction accuracy for a broad range of metabolites (Figure 3a). No kernel improved prediction accuracy of GC-MS metabolites over GBLUP, but the ‘LCGC2’ kernel decreased accuracy in two environments (Figure 3b).

**Figure 3.**
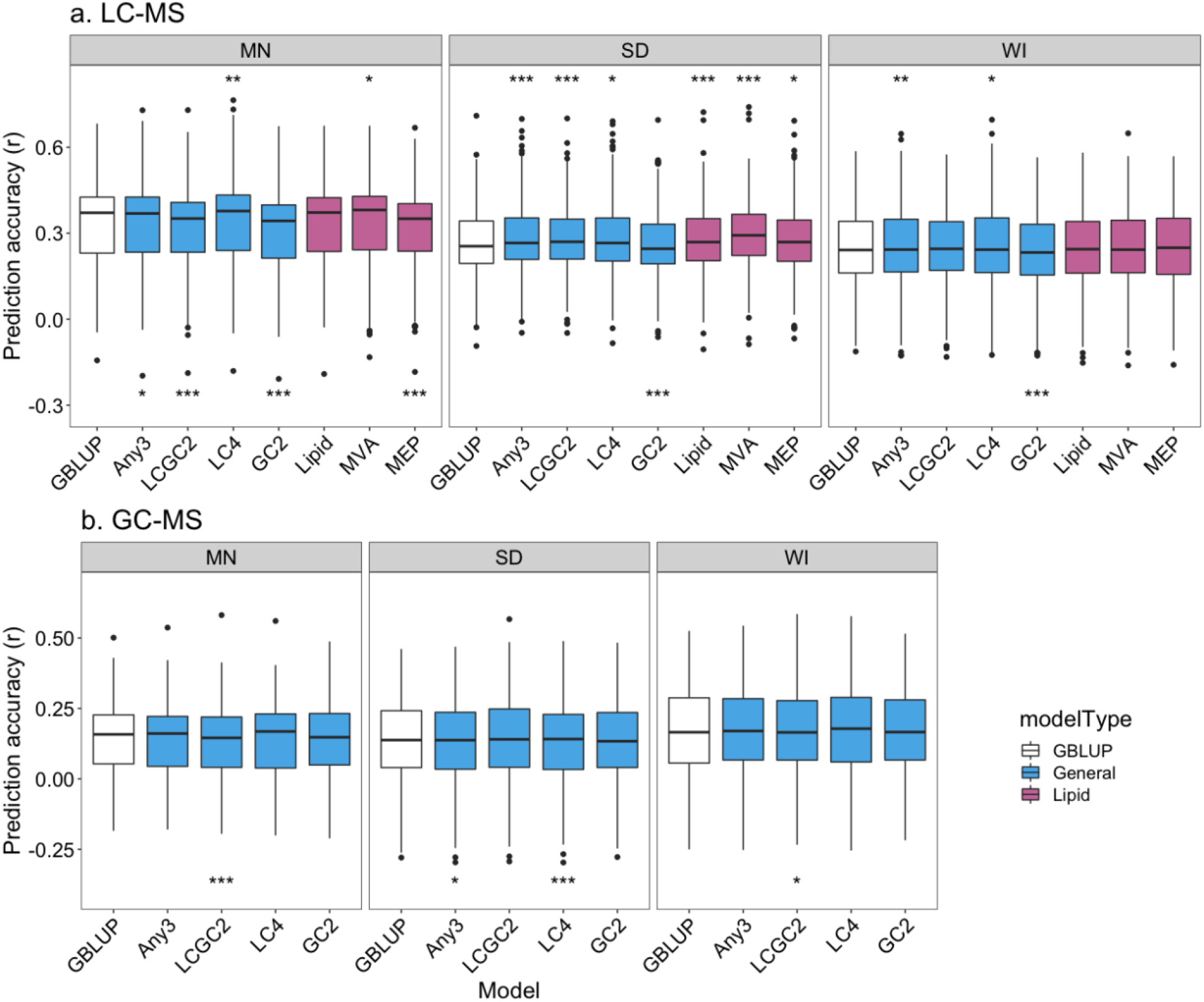
Mean cross-fold validation accuracy (r) of all (a.) LC-MS (*n*=397) and (b.) GC-MS (*n*=243) metabolites by environment (Minnesota, “MN”; South Dakota, “SD” and Wisconsin, “WI”) and two-kernel metabolite model (see Table 2). The models were compared to GBLUP and significant difference indicators are given if the two-kernel metabolite model had higher accuracy than GBLUP at the top of the boxplot, and significance indicators of lower accuracy than GBLUP are given below. The * indicates a *p-*value less than the Bonferroni cutoff per plot, and ** and *** indicate p < 1e-4, and p<1e-6, respectively.

**Table 4.** Mean cross-fold validation accuracy (r) of all LC-MS (*n*=397) and GC-MS (*n*=243) metabolites by (Minnesota, “MN”; South Dakota, “SD” and Wisconsin, “WI”) and model (see Table 2). The color indicates relative value, where blue are highest values and red are lowest values, coded by instrument.

| Environment | Model |  | LCMS | GCMS |
| --- | --- | --- | --- | --- |
| MN | *GBLUP* |  | 0.325 | 0.138 |
|  | General | Any3 | 0.329 | 0.137 |
|  |  | LCGC2 | 0.313 | 0.132 |
|  |  | LC4 | 0.336 | 0.140 |
|  |  | GC2 | 0.303 | 0.137 |
|  | Lipid | Lipid | 0.326 | NA |
|  |  | MEP | 0.314 | NA |
|  |  | MVA | 0.334 | NA |
| SD | *GBLUP* |  | 0.268 | 0.138 |
|  | General | Any3 | 0.280 | 0.135 |
|  |  | LCGC2 | 0.278 | 0.131 |
|  |  | LC4 | 0.278 | 0.141 |
|  |  | GC2 | 0.261 | 0.136 |
|  | Lipid | Lipid | 0.278 | NA |
|  |  | MEP | 0.273 | NA |
|  |  | MVA | 0.296 | NA |
| WI | *GBLUP* |  | 0.249 | 0.167 |
|  | General | Any3 | 0.256 | 0.167 |
|  |  | LCGC2 | 0.251 | 0.163 |
|  |  | LC4 | 0.256 | 0.172 |
|  |  | GC2 | 0.239 | 0.168 |
|  | Lipid | Lipid | 0.248 | NA |
|  |  | MEP | 0.250 | NA |
|  |  | MVA | 0.250 | NA |

Individual metabolites with higher genomic heritability had greater prediction accuracy (R^2^ _adj_ =0.61-0.79; **Figure S9**). Using paired tests to compare the two kernel metabolite models to GBLUP for each metabolite, the most metabolites (LC-MS and GC-MS) with significant improvements in accuracy were for the ‘MVA’, ‘LC4’ and ‘Any3’ kernels, while the most metabolites with significant reductions in accuracy where for ‘GC2’ and ‘MEP’ kernels for LC-MS metabolites, and no clear patterns for GC-MS metabolites (Table 5). On average, 37% and 26% of LC-MS and GC-MS metabolites, respectively had higher prediction accuracy with any of the two-kernel metabolite models than GBLUP, and 47% and 28% had lower prediction accuracy with any of the two-kernel metabolite models than GBLUP. Of all metabolites identified to have significant changes in accuracy compared to GBLUP, two-thirds were unique to one environment (**Figure S10**).

**Table 5.** Number of metabolites (of 397 LC-MS and 243 GC-MS metabolites) where the cross-fold validation accuracy (r) of the given metabolite model (see Table 2) is significantly greater or less than the accuracy of GBLUP. The environments are: Minnesota, “MN”, South Dakota, “SD” and Wisconsin, “WI”. The color indicates relative value, where blue are highest values and red are lowest values, coded by column.

| Type | Model | Env | LCMS |  |  | GCMS |  |
| --- | --- | --- | --- | --- | --- | --- | --- |
|  |  |  | n_better | n_worse |  | n_better | n_worse |
| General | Any3 | MN | 39 | 24 |  | 29 | 40 |
|  |  | SD | 59 | 14 |  | 15 | 31 |
|  |  | WI | 31 | 20 |  | 31 | 23 |
|  | LCGC2 | MN | 7 | 158 |  | 31 | 33 |
|  |  | SD | 61 | 16 |  | 12 | 18 |
|  |  | WI | 36 | 22 |  | 26 | 18 |
|  | LC4 | MN | 41 | 27 |  | 36 | 44 |
|  |  | SD | 55 | 41 |  | 13 | 40 |
|  |  | WI | 30 | 30 |  | 39 | 26 |
|  | GC2 | MN | 6 | 122 |  | 16 | 39 |
|  |  | SD | 9 | 41 |  | 19 | 17 |
|  |  | WI | 8 | 62 |  | 16 | 35 |
| Lipid | Lipid | MN | 20 | 22 |  | NA | NA |
|  |  | SD | 64 | 20 |  | NA | NA |
|  |  | WI | 18 | 27 |  | NA | NA |
|  | MEP | MN | 20 | 104 |  | NA | NA |
|  |  | SD | 53 | 30 |  | NA | NA |
|  |  | WI | 40 | 43 |  | NA | NA |
|  | MVA | MN | 44 | 29 |  | NA | NA |
|  |  | SD | 133 | 20 |  | NA | NA |
|  |  | WI | 32 | 36 |  | NA | NA |

Using the metabolite kernel and the rest-of-genome kernel to partition genetic variation, we found that the metabolite kernels consistently accounted for almost half of total heritability (Figure 4). The ‘Any3’ and ‘LC4’ kernels accounted for more percent heritability for LC-MS than GC-MS metabolites in two environments, and the ‘GC2’ kernel accounted for more percent heritability explained for GC-MS than LC-MS metabolites in all environments (Figure 4a). Percent heritability explained was generally lower in MN than SD and WI for LC-MS metabolites (Figure 4b), and there were differences observed between environments for GC-MS metabolites for the ‘LCGC2’ and ‘GC2’ kernels (Figure 4c). There were weak negative relationships between metabolite genomic heritability and percent heritability explained by the kernel (R^2^ _adj_ =0.05 – 0.15; **Figure S11**), but no relationship between percent heritability explained by the kernel with kernel size (**Table S7**).

**Figure 4.**
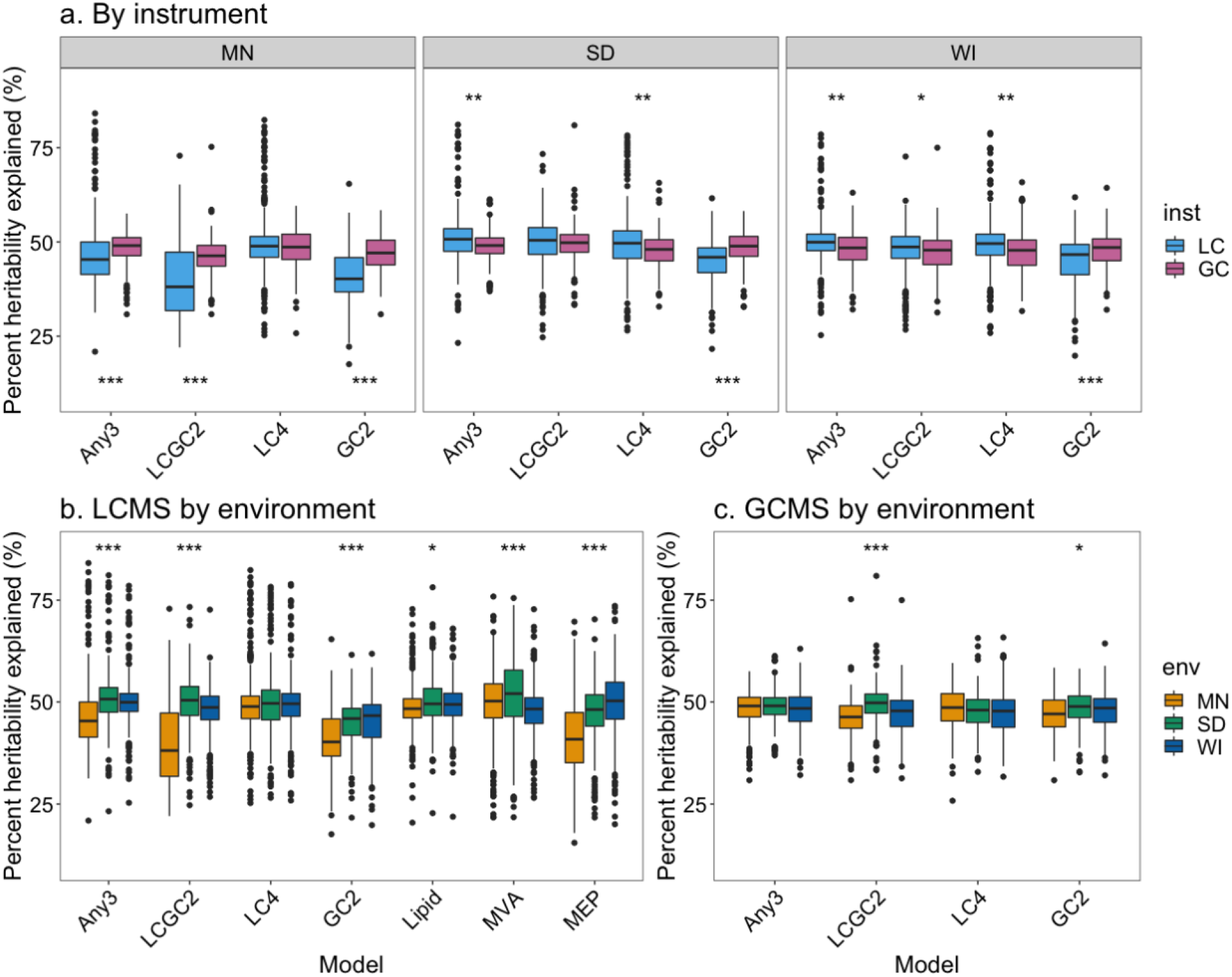
Percent genetic variation attributed to the metabolite kernel for LC-MS (*n*=397) and GC-MS (*n*=243) metabolites in all environments (Minnesota, “MN”; South Dakota, “SD” and Wisconsin, “WI”). (a) The difference in percent genetic variation attributed to metabolite kernel between LC-MS and GC-MS metabolites, where significance indicators above the boxplot represent if percent variation is greater for LC-MS metabolites and below the boxplot if percent variation is greater for GC-MS metabolites. The difference between environments for (b) all metabolite models for LC-MS and (c) all general models for GC-MS instrument. The * indicates a *p-*value less than the Bonferroni cutoff per plot, and ** and *** indicate p < 1e-4, and p<1e-6, respectively.

We compared genetic variation attributed to metabolite kernels to random kernels of similar sizes, constructed from SNPs that were significant (*p_FDR_* < 0.05) mGWAS results that were not included in kernels, and found that, for LC-MS metabolites, the ‘Any3’, ‘LC4’, ‘Lipid’ and ‘MVA’ kernels explained more genetic variance but the ‘LCGC2’ and ‘GC2’ kernels explained less (**Table S7**). In contrast, metabolite kernels never explained more percent genetic variation than random mGWAS kernels for GC-MS metabolites (**Table S7**).

To better understand the effect of the environment on relative model outcomes, we calculated the rank correlation of metabolite prediction accuracy between models and performed hierarchical clustering of the Euclidean distance between ranks. For all metabolites, the ‘Any3’ and ‘LC4’ and the ‘LCGC2’ and ‘GC2’ kernels grouped in all environments (Figure 5).

**Figure 5.**
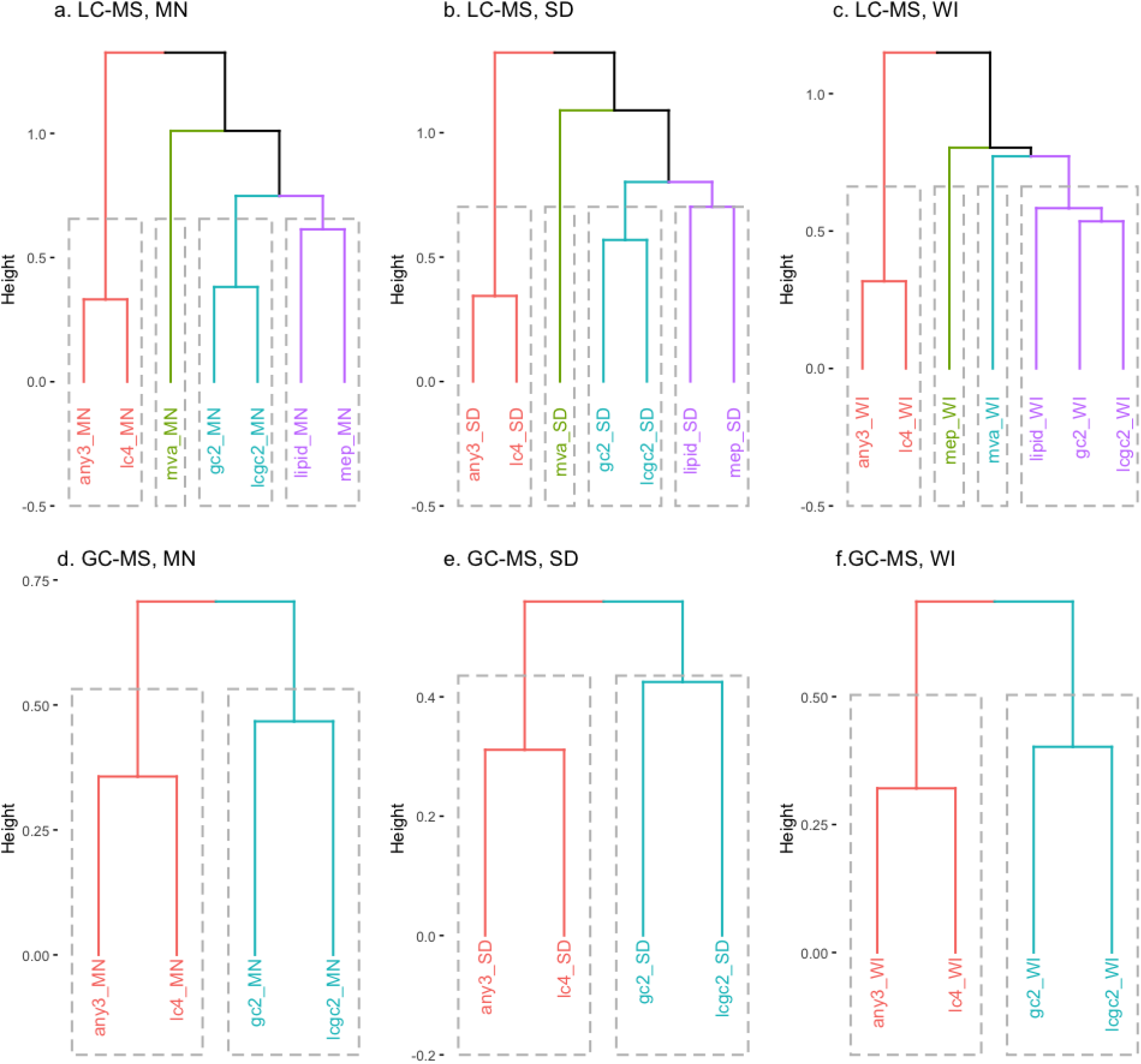
Dendrograms of distance in metabolite kernel performance for (a.-c.) LC-MS (*n*=397) and (d.-f.) GC-MS (*n*=243) metabolites by environment (Minnesota, “MN”; South Dakota, “SD” and Wisconsin, “WI”). Four hierarchical clusters are indicated by color and dashed box.

### Grouping metabolites by similarity

We evaluated if similar metabolites had similar model rankings, where we defined metabolite similarity by: (1) known annotations, (2) structural characteristics as classified by ClassyFire, and (3) Euclidean distance between phenotypes.

For seven oat specialized metabolites where high-confidence named annotations are available (avenanthramides, avenacins, avenacosides), there were 24 instances (of the 147 trait, model and environment combinations) where including a metabolite kernel significantly changed prediction accuracy compared to GBLUP (**Table S8**). We found that similar metabolites had similar ranks of kernels by prediction accuracy in two environments (MN, WI) (Figure 6). These results indicate that when we have access to high-confidence named annotations to define similar metabolites, the similar metabolites have similar prediction results.

**Figure 6.**
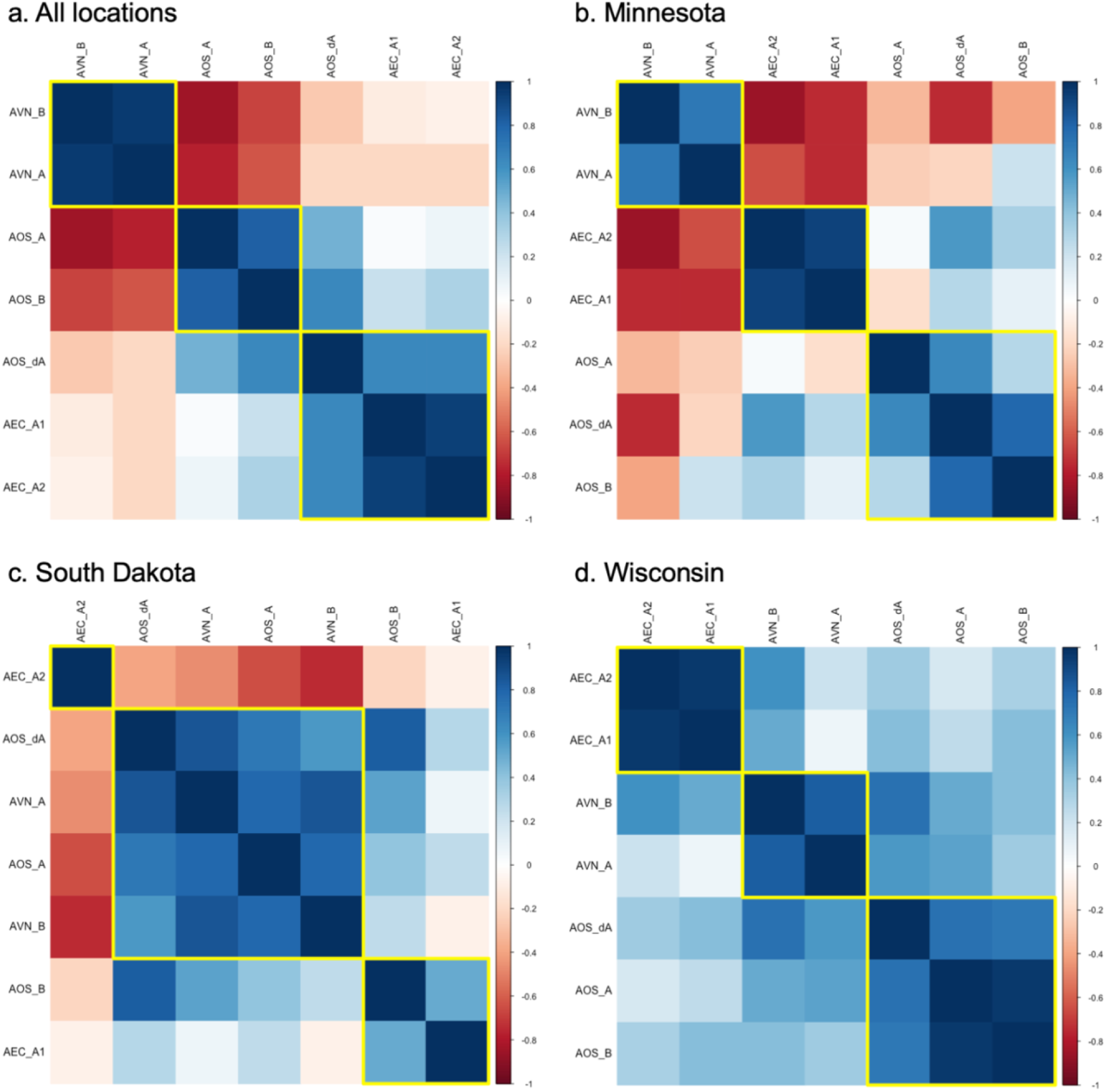
Correlograms of metabolite kernel prediction accuracy rank correlation for seven oat specialized metabolites by (a.) all environments together, and (b.-d.) by individual environment. A color indicator of correlation is shown for all correlations. The yellow boxes represent hierarchical clustering for *n*=3. The metabolite abbreviations are as follows: AVN_A, avenanthramide A; AVN_B, avenanthramide B; AEC_A1, AEC_A2, avenacin A1; AOS_A, avenacoside A; AOS_dA, 26-Desglucoavenacoside A; AOS_B, avenacoside B.

We assessed LC-MS metabolites structurally classified as lipids (*n*=91), and particularly prediction accuracy of the ‘Lipid’ compared to others. While the ‘Lipid’ two-kernel model significantly outperformed GBLUP in only one environment (SD), it generally had higher prediction accuracy than most other kernels besides ‘MVA’ in two environments (MN, SD) (Figure 7). Other kernels accounted for more heritability than the lipid kernel in only two instances (Figure 7). We defined lipids as ‘similar’ by ‘Class’ descriptor (e.g. steroids, or fatty acyls), and anticipated similar model rankings by lipid class. We found lipid Class was not predictive of the model rank (Figure 8), suggesting that structural classifications may not provide effective metabolite groupings.

**Figure 7.**
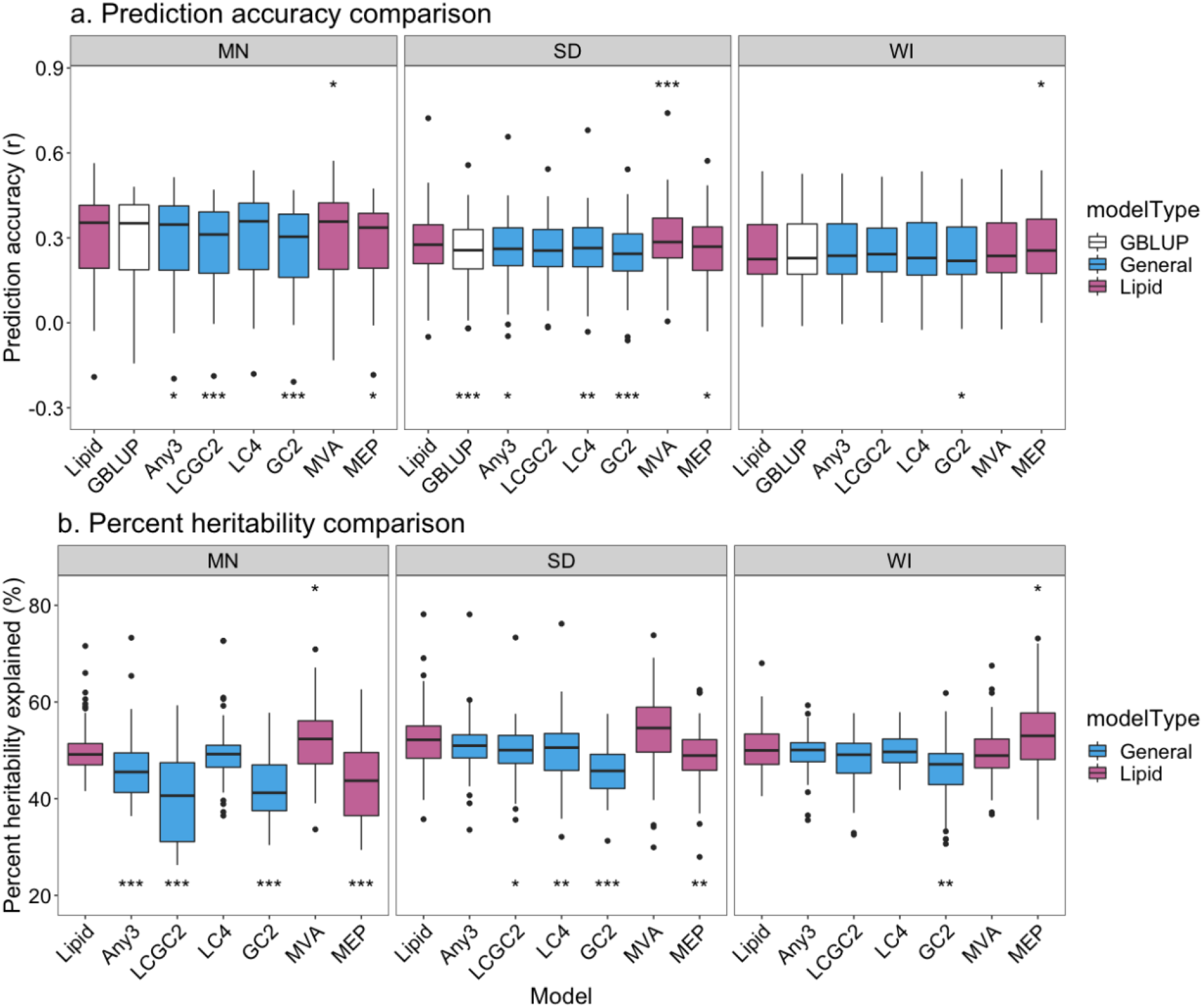
(a) Mean cross-fold validation accuracy (r) of, and (b) percent heritability (genetic variation) attributed to the metabolite kernel for, LC-MS lipid metabolites (*n*=91) by environment (Minnesota, “MN”; South Dakota, “SD” and Wisconsin, “WI”) and two-kernel metabolite model (see Table 2). The models were compared to ‘Lipid’ kernel and significant difference indicators are given if the two-kernel metabolite model had higher accuracy than ‘Lipid’ at the top of the boxplot, and significance indicators of lower accuracy than ‘Lipid’ are given below. The * indicates a *p-*value less than the Bonferroni cutoff per plot, and ** and *** indicate p < 1e-4, and p<1e-6, respectively.

**Figure 8.**
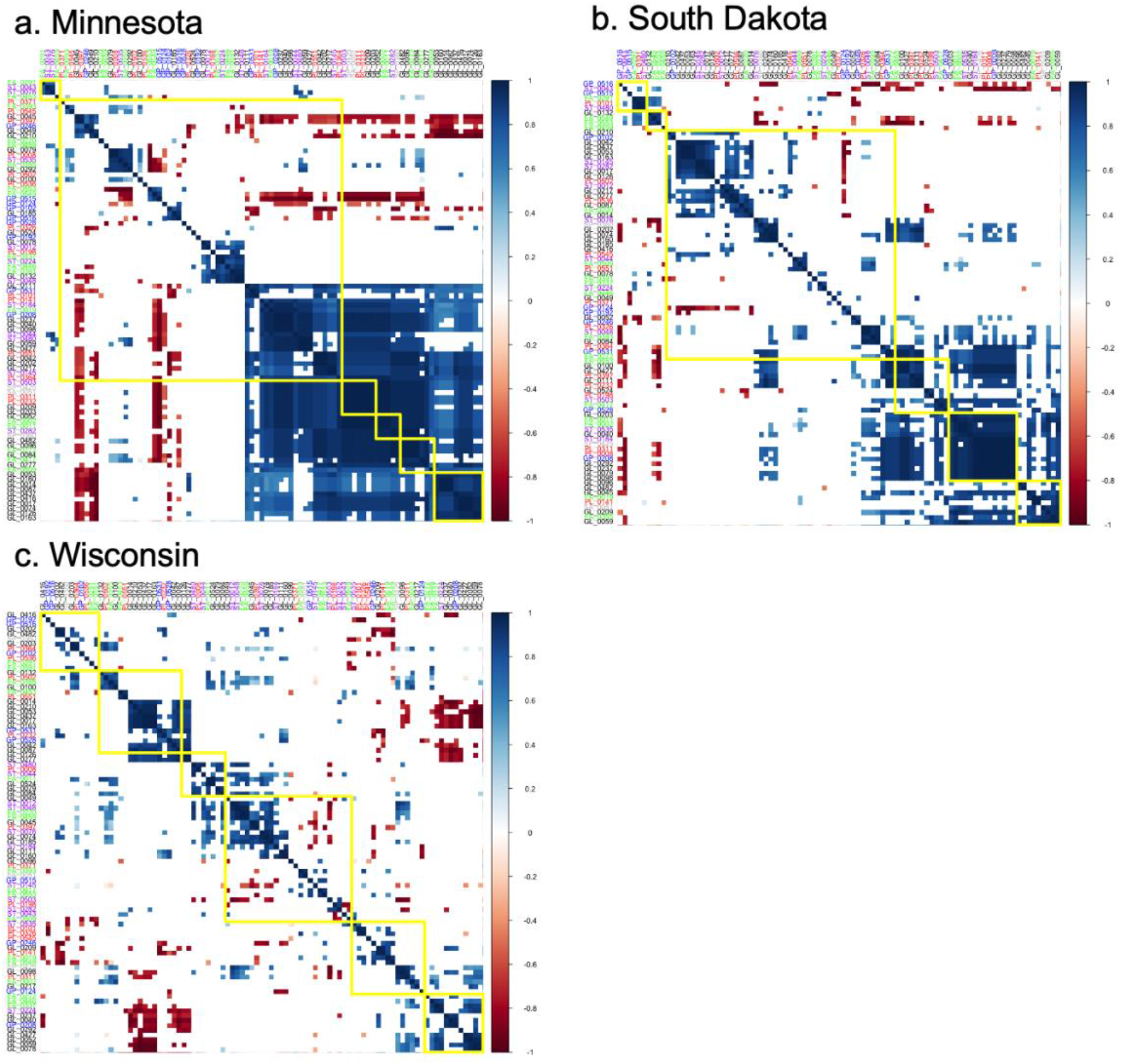
Correlograms of metabolite kernel prediction accuracy rank correlation by model for *n*=91 LC-MS lipids by (a.) all environments together, and by individual environment (b.-d.). A color indicator of correlation is shown for all correlations with *p*<0.05. The text label color indicates type of lipid. The yellow boxes represent hierarchical clustering for *n*=6. The name and color key for lipid type is given in **Table S9**.

Finally, without using annotations, we computed the distance between metabolites and performed hierarchical clustering to define 10 metabolite groups per environment. Most of the groups had significantly higher correlations of model rank within group compared to metabolites out of the group (Figure 9). We found that the groups were largely defined by retention time. Groups with strong within-group correlation had smaller coefficients of variation in retention time (CV<20) than other groups, but the trends in genomic heritability were not consistent between groups (Table 6). These groups also had less variation in retention time than the lipid Classes (CV> 20; **Table S9**).

**Figure 9.**
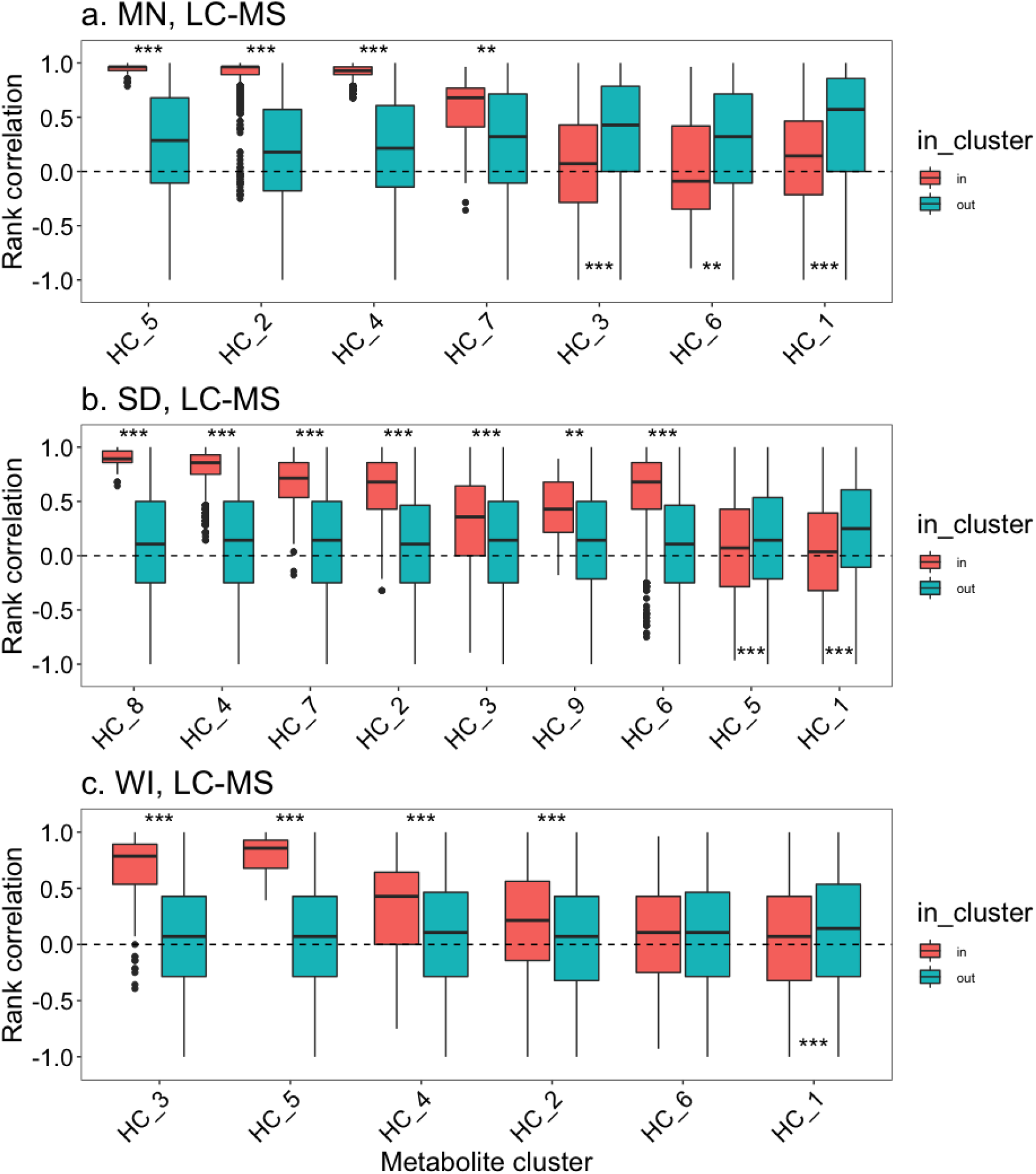
Rank correlation of metabolite kernel prediction accuracy rank correlation by model for groups of LC-MS metabolites defined by hierarchical clustering of a distance metric by environment (Minnesota, “MN”; South Dakota, “SD” and Wisconsin, “WI”). There is no relationship between cluster names across environments. Clusters with 10 or more metabolites are presented with the metabolites within the cluster are shown in blue, and the metabolites not in the cluster are shown in red, and comparisons are made between the two sets by group. Significant difference indicators are given at the top of the boxplot if the metabolites within the group had stronger correlation than those not in the group, and vice versa for significance indicators below. The * indicates a *p-*value less than the Bonferroni cutoff per plot, and ** and *** indicate p < 1e-4, and p<1e-6, respectively.

**Table 6.** Coefficient of variation (“CV”) in retention time (s) and genomic heritability (mean +/- one standard deviation) of LC-MS metabolites by metabolite group defined by hierarchical cluster. Note that there is no relationship between cluster name across environments. The number of metabolites in each group is given by ‘n’. Metabolite groups with ten or more metabolites that had higher within group correlation are indicated with a *.

| Group | MN |  |  |  | SD |  |  |  | WI |  |  |  |
| --- | --- | --- | --- | --- | --- | --- | --- | --- | --- | --- | --- | --- |
|  | *n* | *RT-CV* | *h*<sup>2</sup> |  | *n* | *RT-CV* | *h*<sup>2</sup> |  | *n* | *RT-CV* | *h*<sup>2</sup> |  |
| 1 | 134 | 80.2 | 0.30 +/- 0.19 |  | 123 | 87.7 | 0.22 +/- 0.19 |  | 206 | 70.5 | 0.20 +/- 0.15 |  |
| 2 | 90 | 11.2 | 0.39 +/- 0.07 | * | 29 | 2.8 | 0.09 +/- 0.06 | * | 84 | 13.4 | 0.11 +/- 0.06 | * |
| 3 | 69 | 24.2 | 0.17 +/- 0.12 |  | 44 | 21.8 | 0.24 +/- 0.17 | * | 28 | 2.7 | 0.11 +/- 0.05 | * |
| 4 | 54 | 3.2 | 0.42 +/- 0.04 | * | 47 | 2.7 | 0.10 +/- 0.03 | * | 26 | 3.2 | 0.08 +/- 0.04 | * |
| 5 | 16 | 1.4 | 0.40 +/- 0.02 | * | 52 | 25.7 | 0.21 +/- 0.15 |  | 11 | 6.6 | 0.39 +/- 0.07 | * |
| 6 | 12 | 10.1 | 0.08 +/- 0.07 |  | 54 | 15.8 | 0.12 +/- 0.06 | * | 31 | 24.7 | 0.18 +/- 0.14 |  |
| 7 | 11 | 6.3 | 0.10 +/- 0.06 | * | 21 | 2.0 | 0.07 +/- 0.04 | * | 4 | 1.3 | 0.22 +/- 0.03 | NA |
| 8 | 5 | 9.6 | 0.18 +/- 0.16 | NA | 11 | 6.6 | 0.26 +/- 0.04 | * | 3 | 7.4 | 0.20 +/- 0.06 | NA |
| 9 | 3 | 7.4 | 0.14 +/- 0.08 | NA | 10 | 6.5 | 0.10 +/- 0.04 | * | 2 | 2.8 | 0.48 +/- 0.13 | NA |
| 10 | 2 | 2.8 | 0.47 +/- 0.07 | NA | 5 | 1.3 | 0.19 +/- 0.06 | NA | 1 | NA | NA | NA |

## DISCUSSION

Our work tests generalizable frameworks for genomic prediction of a diverse array of plant metabolites. Using a discovery germplasm panel, we identified loci by mGWAS that represent different biological bases – loci that affect multiple types of metabolites to metabolites from specific biochemical pathways. Building kernels from significant mGWAS loci that affect multiple LC-MS metabolites and specific pathways thereof increased prediction accuracy over GBLUP in a validation panel for LC-MS metabolites. No model tested improved prediction of GC-MS metabolites over GBLUP, and kernels from GC-MS metabolites reduced prediction accuracy in some cases. mGWAS-defined kernels accounted for ∼45% of genetic variation, and rank of kernel performance was consistent between environments. An ongoing challenge in developing generalized genomic prediction frameworks is defining metabolite ‘similarity’. We found that grouping metabolites by high-confidence named annotations and computationally derived groupings (without annotations) had similar outcomes from the models tested, while metabolites delineated by structural features alone did not. Overall, this work builds from efforts to predict tens of biochemically similar metabolites to metabolome-wide genomic prediction.

### Characterizing the oat metabolome by mGWAS

We evaluated over 2000 metabolites measured by LC-MS or GC-MS in mature oat seed and found that, on average, metabolites had low to moderate genomic heritability (mean *h^2^*=0.09 to 0.30), with LC-MS metabolites being more heritable than GC-MS metabolites. Other analyses of untargeted metabolites (*n*=900-7000 metabolites) report wide ranges of broad-sense (not genomic) heritability (H^2^), from a uniform distribution (Zhou et al., 2019), to right (Zhu et al., 2018) and left (Chen et al., 2016) skews. While some differences in heritability between studies could be attributed to the tissue and developmental specificity of metabolites (Soltis & Kliebenstein, 2015), we also found that metabolite heritability covaries with column retention time (related to metabolite polarity). While retention time was not evaluated, (Zhou et al., 2019) found that less common features tended to have lower heritability that they attributed to machine artifact. This suggests that parameters such as specific extraction (e.g., if the extracting solvent more efficiently extracts polar or non-polar compounds), or signal processing methods may affect error variation.

By conducting mGWAS for the 1668 metabolites in the discovery panel, we found that a greater proportion of LC-MS than GC-MS metabolites had significant mGWAS results, even when controlling for heritability differences, suggesting that more LC-MS metabolites have an oligogenic genetic architecture. Overall, primary metabolites (measured by GC-MS) tend to be dominantly inherited (Schauer et al., 2008; Fernie & Tohge, 2017), and variation is determined by multiple small effect loci (Soltis & Kliebenstein, 2015). In contrast, specialized metabolites (measured by LC-MS) generally arise from variation in primary metabolism (Moghe & Last, 2015; Maeda, 2019) including enzyme neofunctionalization (Pichersky & Gang, 2000; Fernie & Tohge, 2017). Nonetheless, selection type (e.g., direction or stabilizing) in crops is more important in predicting loci effects than type of metabolite *per se* (Soltis & Kliebenstein, 2015). There are multiple examples of balancing selection for metabolite concentration (e.g., as defensive metabolite, or regionally preferred crop aesthetic or flavor) (Soltis & Kliebenstein, 2015), and (Campbell et al., 2021b) proposed that optimizing or stabilizing selection pressures predominately shape the oat seed metabolome.

Another factor that may contribute to the differences between the mGWAS results for GC-MS (primary) and LC-MS (specialized) metabolites is that metabolites were measured in mature seed. Primary metabolites decreased in *Arabidopsis* seed during reserve accumulation, but then increased during seed desiccation (putatively for availability for germination energy) (Fait et al., 2006). In contrast, primary metabolites in rice consistently decrease beginning at desiccation (Hu et al., 2016). In a time-series transcriptome-wide analysis of developing oat seed, expressed genes had enriched GO terms for photosynthesis until 23 days after anthesis (DAA), followed by an enrichment of GO terms for nutrient reservoir activity beginning at 28 DAA (Hu et al., 2020). These results suggest that the metabolomic dynamics in developing oat seed may be similar to those of rice, and point to a need for multiple metabolome measures during seed development.

### Potential for generalizable approaches for genomic prediction of metabolites

We used multiple criteria for constructing metabolite kernels to test hypotheses of which biological partition may be the most enriched for causal SNPs. We developed kernels to encompass general metabolome-wide information from both or single LC-MS and GC-MS instruments (‘Any3’, ‘LCGC2’, and LC4’, ‘GC2’, respectively), or metabolites structurally identified as lipids (‘Lipid’), and pathways thereof (‘MVA’, ‘MEP’) for two-kernel genomic prediction. Importantly, to make our results relevant for plant breeding programs, we selected SNPs from a diverse ‘discovery’ panel, and evaluated prediction accuracy in another more elite population evaluated in multiple environments.

Metabolite kernels accounted for a high percent of trait genetic variation, and the ‘Any3’, ‘LC4’, and ‘MVA’ kernels consistently increased prediction accuracy over GBLUP for LC-MS metabolites. While the ‘MVA’ kernels included the highest number of SNPs in genes of any of the kernels, high gene richness did not always translate to high prediction accuracy (e.g., the ‘Lipid’ kernel), indicating that gene richness alone does not account for our results.

The general kernels likely include loci that affect multiple metabolites, as loci with pleiotropy and epistatic interactions are common for metabolites (Soltis & Kliebenstein, 2015), and we hypothesized that using these kernels would increase prediction accuracy of the most metabolites. The ‘Any3’ and ‘LC4’ kernels improved prediction accuracy, and the ‘LC4’ kernel more so, where the ‘LC4’ kernel is a subset of the ‘Any3’ kernel. Our approach can be compared to factor analysis recently used in genomic prediction of several oat fatty acids (Campbell et al., 2021b). In both cases (results from individual mGWAS and result from GWAS of factors), multi-kernel models improved prediction accuracy. Nonetheless, many factors extracted from oat metabolomic data were enriched for lipids (Campbell et al., 2021b), while our ‘Any3’ and ‘LC4’ kernels were depleted for lipids, indicating that we are capturing different information than the factor analysis. Overall, these results suggest that distilling results from the entire metabolome identifies SNPs that affect multiple metabolites and improves prediction accuracy.

Contrary to our expectations, the ‘MVA’ kernel that incorporated only a specific branch of terpenoid biosynthesis (e.g., triterpenoids and sesquiterpenoids) improved prediction accuracy of LC-MS metabolites metabolome-wide as much as the general ‘LC4’ and ‘Any3’ kernels. While the ‘MEP’ kernel representing another terpenoid biosynthetic pathway (e.g., diterpenoids and carotenoids) did not improve accuracy, these pathways function largely independently, and sometimes antagonistically (Rodríguez-Concepción & Boronat, 2015). Increased prediction accuracy from the ‘MVA’ kernel suggests that loci governing variation in specific pathways may translate across populations for metabolome-wide prediction. Alternatively, this result could be specific to terpenoids: (Turner-Hissong et al., 2020) reported that a terpenoid gene kernel improved prediction of a free amino acid, isoleucine, in *Arabidopsis* seed where the terpenoids are unrelated to isoleucine biosynthesis. It would be intriguing to test if terpenoid-related kernels improve prediction accuracy of seemingly unrelated metabolites in other non-seed tissues (with lower oil content) to assess if energetic tradeoffs are responsible for this observation.

A kernel derived from mGWAS results from LC-MS metabolites structurally identified as lipids (‘Lipid’) in the discovery panel, did not improve prediction accuracy metabolome-wide, or for lipids over GBLUP in the validation panel. (Campbell et al., 2021b) found that latent factors that were enriched for lipids did not significantly improve prediction accuracy of proteins, likely due to high negative genetic correlation between those traits and that factor loadings included more metabolites than just lipids. The ‘Lipid’ kernel here was also potentially too expansive of a categorization and may have led to kernels containing genomic regions with shared regulation but opposing effects. This result suggests that grouping metabolites by shared regulatory control may be more beneficial (e.g., ‘MVA’), and will become more feasible with improved genomic resources.

Finally, no method we tested improved prediction accuracy of GC-MS metabolites, and kernels from solely mGWAS results from GC-MS metabolites (‘GC2’) *reduced* prediction accuracy of LC-MS metabolites. This may be because GC-MS metabolites had lower heritability (potentially due to lower phenotypic variation in mature seed, constraints on potential genetic variation), fewer mGWAS results, and thus provided less reliable information. Overall, these results highlight that combining multiple metabolomics datasets from different instruments may have limited efficacy, depending on, for instance, development stage sampled.

### Strategies for categorizing ‘similar’ metabolites

In building generalized frameworks, it would be useful to have high-throughput methods for identifying similar metabolites to which to apply the same prediction method. A key challenge, however, is how ‘similar’ is defined. We tested three definitions of ‘similar’: high-confidence named annotations of known metabolites (difficult to obtain, high biological information), automated metabolite classification by chemical structure (moderate effort to obtain, some biological information), and by an annotation-free measure of similarity (easy to obtain, no biological information). Overall, groups of metabolites by named annotations and by the annotation-free measure, had consistent ranks of the models tested. In the annotation-free grouping, we found that retention time was an important predictor of group association. As metabolite annotations provide useful biological information, we look forward to more high confidence annotations as databases continue to grow (Afendi et al., 2012).

Defining ‘similar’ by structural classification was the least successful method, perhaps because structural classifications do not broadly correspond to a biosynthetic pathway (Djoumbou Feunang et al., 2016). A caveat in examining relative model rankings is that we did not specifically design kernels to evenly represent the space of all potential kernels but, as the purpose of this study was to test different biological rationales, this analysis is informative for understanding differences between approaches.

## CONCLUSIONS

We are building towards a generalized framework for genomic prediction of metabolites by investigating how we can efficiently extract information from metabolomics data, integrate biology to find the most informative loci, and then test for which metabolites these strategies are most successful. Our work extends the foundational metabolomics work done in model organisms like *Arabidopsis*, tomato, maize (Fernie & Tohge, 2017) and on conserved biochemical pathways (Wager & Li, 2018), to provide strategies for genomic prediction of multiple, diverse metabolites in non-model crops. Overall, we show that integrating whole metabolome or specific pathway information improves genomic prediction accuracy and translates across populations within a species. This work also provides a framework for testing such models between closely related species by transfer learning (Wang et al., 2020).

## Abbreviations

drBLUPs: deregressed best linear unbiased predictors
GC-MS: gas chromatography - mass spectrometry
LC-MS: liquid chromatography - mass spectrometry
MEP: Methylerythritol Phosphate pathway
mGWAS: metabolite genome wide association study
MVA: Mevalonate Acid pathway

## ACKNOWLEDGEMENTS

Corey D. Broeckling conducted the metabolomics extraction, measurement, and data processing at the Bioanalysis and Omics Center of the Analytical Resources Core at Colorado State University (Fort Collins, CO USA). Funding for this research was provided by the United States Department of Agriculture - National Institute of Food and Agriculture - Agriculture and Food Research Initiative (USDA NIFA-AFRI) grant (2017-67007-26502), and the USDA Agricultural Research Service Project Number 8062-21000-045-000-D.

## AUTHOR CONTRIBUTIONS

JLJ, MAG and MES designed the research. LJB, HH, and MTC analyzed the data, and HH, MTC, MC, LG, KPS and MES conducted experiments. LJB, MAG and JLJ wrote the manuscript and all co-authors were involved in editing the manuscript.

## DATA AVAILABILITY

Deregressed BLUPs of the metabolites for the discovery (diverse) germplasm panel are available in the supplementary material of (Campbell et al., 2021b). Deregressed BLUPs of the metabolites for the validation germplasm panel is provided as **Supporting File 1**. Genotype data is as used in (Brzozowski et al., 2021) and available at https://datacommons.cyverse.org/browse/iplant/home/shared/GoreLab/dataFromPubs/Brzozowski_OatMetabolome_2021. The R code for these analyses is available on a public repository in https://github.com/ljbrzozowski/OatMetaboliteGenomicPrediction

## SUPPORTING INFORMATION CONTENTS

**File S1**. Validation germplasm panel metabolite information

**Figure S1.** Linear regressions of metabolite heritability by retention time and molecular mass.

**Figure S2.** Number of metabolites with significant GWAS results by instrument type.

**Figure S3.** The number of metabolites and SNPs contributing to kernels.

**Figure S4.** Correlation between off-diagonal elements in metabolite kernels.

**Figure S5.** Mean Euclidean distance between metabolites that contribute to metabolite kernels.

**Figure S6.** Genomic distribution of metabolite GWAS results and the general kernels.

**Figure S7.** Genomic distribution of metabolite GWAS results and the lipid kernels.

**Figure S8.** Validation panel metabolite genomic heritability and relationship to retention time.

**Figure S9.** Mean cross-fold validation accuracy of metabolites compared to genomic heritability

**Figure S10.** Metabolites where any two-kernel metabolite model improved or reduced genomic prediction accuracy over GBLUP.

**Figure S11.** Percent genetic variation attributed to the metabolite kernel compared to metabolite genomic heritability.

**Table S1.** Mean metabolite heritability by ClassyFire superclass groups for the discovery panel.

**Table S2.** Number of metabolites in each metabolite kernel.

**Table S3.** Number of SNPs in each metabolite kernel.

**Table S4.** Spearman’s rank correlation of metabolite heritability across validation panel environments.

**Table S5.** Validation panel LC-MS metabolite ClassyFire ‘Superclass’ count and heritability.

**Table S6.** Validation panel linear regression between retention time and heritability

**Table S7.** Comparison of percent genetic variance (percent heritability) by metabolite kernels to mGWAS random kernels by instrument, model and environment.

**Table S8.** Significant differences in prediction accuracy compared to GBLUP for seven specialized metabolites.

**Table S9.** Coefficient of variation in retention time of LC-MS lipids by Class.

## REFERENCES

Afendi, F.M., Okada, T., Yamazaki, M., Hirai-Morita, A., Nakamura, Y., Nakamura, K., Ikeda, S., Takahashi, H., Altaf-Ul-Amin, M., Darusman, L.K., Saito, K., & Kanaya, S. (2012). KNApSAcK family databases: Integrated metabolite-plant species databases for multifaceted plant research. Plant and Cell Physiology, 53, 1–12. https://doi.org/10.1093/pcp/pcr165

Baseggio, M., Murray, M., Magallanes-Lundback, M., Kaczmar, N., Chamness, J., Buckler, E.S., Smith, M.E., DellaPenna, D., Tracy, W.F., & Gore, M.A. (2019). Genome-Wide Association and Genomic Prediction Models of Tocochromanols in Fresh Sweet Corn Kernels. The Plant Genome, 12, 180038. https://doi.org/10.3835/plantgenome2018.06.0038

Broeckling, C.D., Ganna, A., Layer, M., Brown, K., Sutton, B., Ingelsson, E., Peers, G., & Prenni, J.E. (2016). Enabling Efficient and Confident Annotation of LC−MS Metabolomics Data through MS1 Spectrum and Time Prediction. Analytical Chemistry, 88, 9226–9234. https://doi.org/10.1021/acs.analchem.6b02479

Brzozowski, L.J., Hu, H., Campbell, M.T., Broeckling, C.D., Caffe-Treml, M., Gutiérrez, L., Smith, K.P., Sorrells, M.E., Gore, M.A., & Jannink, J.-L. (2021). Selection for seed size has indirectly shaped specialized metabolite abundance in oat (*Avena sativa* L.). BioRvix,. https://doi.org/10.1101/2021.08.18.454785

Campbell, M.T., Hu, H., Yeats, T.H., Brzozowski, L.J., Caffe-Treml, M., Gutiérrez, L., Smith, K.P., Sorrells, M.E., Gore, M.A., & Jannink, J.-L. (2021)(a). Improving Genomic Prediction for Seed Quality Traits in Oat (*Avena sativa* L.) Using Trait-Specific Relationship Matrices. Frontiers in Genetics, 12, 643733. https://doi.org/10.3389/fgene.2021.643733

Campbell, M.T., Hu, H., Yeats, T.H., Caffe-Treml, M., Gutiérrez, L., Smith, K.P., Sorrells, M.E., Gore, M.A., & Jannink, J.-L. (2021)(b). Translating insights from the seed metabolome into improved prediction for lipid-composition traits in oat (*Avena sativa* L.). Genetics, 217, iyaa043. https://doi.org/10.1093/genetics/iyaa043

de los Campos, G., Sorensen, D., & Gianola, D. (2015). Genomic Heritability: What Is It?. PLOS Genetics, 11, e1005048. https://doi.org/10.1371/journal.pgen.1005048

Carlson, M.O., Montilla-Bascon, G., Hoekenga, O.A., Tinker, N.A., Poland, J., Baseggio, M., Sorrells, M.E., Jannink, J.L., Gore, M.A., & Yeats, T.H. (2019). Multivariate genome-wide association analyses reveal the genetic basis of seed fatty acid composition in oat (*Avena sativa* L.). G3: Genes, Genomes, Genetics, 9, 2963–2975. https://doi.org/10.1534/g3.119.400228

Chen, W., Wang, W., Peng, M., Gong, L., Gao, Y., Wan, J., Wang, S., Shi, L., Zhou, B., Li, Z., Peng, X., Yang, C., Qu, L., Liu, X., & Luo, J. (2016). Comparative and parallel genome-wide association studies for metabolic and agronomic traits in cereals. Nature Communications, 7, 12767. https://doi.org/10.1038/ncomms12767

Cuevas, J., Montesinos-López, O.A., Martini, J.W.R., Pérez-Rodríguez, P., Lillemo, M., & Crossa, J. (2020). Approximate Genome-Based Kernel Models for Large Data Sets Including Main Effects and Interactions. Frontiers in Genetics, 11, 567757. https://doi.org/10.3389/fgene.2020.567757

Diepenbrock, C.H., Ilut, D.C., Magallanes-Lundback, M., Kandianis, C.B., Lipka, A.E., Bradbury, P.J., Holland, J.B., Hamilton, J.P., Wooldridge, E., Vaillancourt, B., Góngora-Castillo, E., Wallace, J.G., Cepela, J., Mateos-Hernandez, M., Owens, B.F., Tiede, T., Buckler, E.S., Rocheford, T., Buell, C.R., Gore, M.A., & DellaPenna, D. (2021). Eleven biosynthetic genes explain the majority of natural variation in carotenoid levels in maize grain. The Plant Cell, 33, 882–900. https://doi.org/10.1093/plcell/koab032

Diepenbrock, C.H., Kandianis, C.B., Lipka, A.E., Magallanes-Lundback, M., Vaillancourt, B., Góngora-Castillo, E., Wallace, J.G., Cepela, J., Mesberg, A., Bradbury, P.J., Ilut, D.C., Mateos-Hernandez, M., Hamilton, J., Owens, B.F., Tiede, T., Buckler, E.S., Rocheford, T., Buell, C.R., Gore, M.A., & DellaPenna, D. (2017). Novel Loci Underlie Natural Variation in Vitamin E Levels in Maize Grain. The Plant Cell, 29, 2374–2392. https://doi.org/10.1105/tpc.17.00475

Djoumbou Feunang, Y., Eisner, R., Knox, C., Chepelev, L., Hastings, J., Owen, G., Fahy, E., Steinbeck, C., Subramanian, S., Bolton, E., Greiner, R., & Wishart, D.S. (2016). ClassyFire: automated chemical classification with a comprehensive, computable taxonomy. Journal of Cheminformatics, 8, 1–20. https://doi.org/10.1186/s13321-016-0174-y

Endelman, J.B. (2011). Ridge Regression and Other Kernels for Genomic Selection with R Package rrBLUP. The Plant Genome, 4, 250–255. https://doi.org/10.3835/plantgenome2011.08.0024

Fait, A., Angelovici, R., Less, H., Ohad, I., Urbanczyk-Wochniak, E., Fernie, A.R., & Galili, G. (2006). Arabidopsis Seed Development and Germination Is Associated with Temporally Distinct Metabolic Switches. Plant Physiology, 142, 839–854. https://doi.org/10.1104/pp.106.086694

Fernie, A.R., & Tohge, T. (2017). The Genetics of Plant Metabolism. Annual Review of Genetics, 51, 287–310

Friedman, J., Hastie, T., & Tibshirani, R. (2010). Regularization Paths for Generalized Linear Models via Coordinate DescentJournal of. Journal of Statistical Software, 33, 1–22

Heffner, E.L., Sorrells, M.E., & Jannink, J.L. (2009). Genomic selection for crop improvement. Crop Science, 49, 1–12. https://doi.org/10.2135/cropsci2008.08.0512

Hernandez, C., Wyatt, L.E., & Mazourek, M. (2020). Genomic Prediction and Selection for Fruit Traits in Winter Squash. G3: Genes, Genomes, Genetics, 10, 3601–3610

Heslot, N., Jannink, J.-L., & Sorrells, M.E. (2015). Perspectives for Genomic Selection Applications and Research in Plants. Crop Science, 55, 1–12. https://doi.org/10.2135/cropsci2014.03.0249

Hu, C., Tohge, T., Chan, S.-A., Song, Y., Rao, J., Cui, B., Lin, H., Wang, L., Fernie, A.R., Zhang, D., & Shi, J. (2016). Identification of Conserved and Diverse Metabolic Shifts during Rice Grain Development. Scientific Reports, 6, 20942. https://doi.org/10.1038/srep20942

Hu, H., Campbell, M.T., Yeats, T.H., Zheng, X., Runcie, D.E., Covarrubias-Pazaran, G., Broeckling, C., Yao, L., Caffe-Treml, M., Gutiérrez, L., Smith, K.P., Tanaka, J., Hoekenga, O.A., Sorrells, M.E., Gore, M.A., & Jannink, J.-L. (2021). Multi-omics prediction of oat agronomic and seed nutritional traits across environments and in distantly related populations. Theoretical and Applied Genetics, *In press*. https://doi.org/10.1101/2021.05.03.442386

Hu, H., Gutierrez-Gonzalez, J.J., Liu, X., Yeats, T.H., Garvin, D.F., Hoekenga, O.A., Sorrells, M.E., Gore, M.A., & Jannink, J.L. (2020). Heritable temporal gene expression patterns correlate with metabolomic seed content in developing hexaploid oat seed. Plant Biotechnology Journal, 18, 1211–1222. https://doi.org/10.1111/pbi.13286

Kumar, R., Bohra, A., Pandey, A.K., Pandey, M.K., & Kumar, A. (2017). Metabolomics for Plant Improvement: Status and Prospects. Frontiers in Plant Science, 8, 1302. https://doi.org/10.3389/fpls.2017.01302

de Los Campos, G. (2018). Various Ways of fitting a “GBLUP” model using BGLR. GitHub,

Luo, J. (2015). Metabolite-based genome-wide association studies in plants. Current Opinion in Plant Biology, 24, 31–38. https://doi.org/10.1016/j.pbi.2015.01.006

Maeda, H.A. (2019). Evolutionary diversification of primary metabolism and its contribution to plant chemical diversity. Frontiers in Plant Science, 10, 1–8. https://doi.org/10.3389/fpls.2019.00881

Moghe, G., & Last, R.L. (2015). Something old, something new: Conserved enzymes and the evolution of novelty in plant specialized metabolism. Plant Physiology, 169, 1512–1523. https://doi.org/10.1104/pp.15.00994

Owens, B.F., Lipka, A.E., Magallanes-Lundback, M., Tiede, T., Diepenbrock, C.H., Kandianis, C.B., Kim, E., Cepela, J., Mateos-Hernandez, M., Buell, C.R., Buckler, E.S., DellaPenna, D., Gore, M.A., & Rocheford, T. (2014). A Foundation for Provitamin A Biofortification of Maize: Genome-Wide Association and Genomic Prediction Models of Carotenoid Levels. Genetics, 198, 1699–1716. https://doi.org/10.1534/genetics.114.169979

Pérez, P., & de los Campos, G. (2014). Genome-Wide Regression and Prediction with the BGLR Statistical Package. Genetics, 198, 483–495. https://doi.org/10.1534/genetics.114.164442

Pichersky, E., & Gang, D.R. (2000). Genetics and biochemistry of secondary metabolites in plants: an evolutionary perspective. Trends in Plant Science, 5, 439–445. https://doi.org/10.1016/S1360-1385(00)01741-6

R Core Team. (2016). R: A Language and Environment for Statistical Computing. R Foundation for Statistical Computing, Vienna, Austria.

Rodríguez-Concepción, M., & Boronat, A. (2015). Breaking new ground in the regulation of the early steps of plant isoprenoid biosynthesis. Current Opinion in Plant Biology, 25, 17–22. https://doi.org/10.1016/j.pbi.2015.04.001

Rossum, B.-J. van, & Kruijer, W. (2020). Package ‘StatgenGWAS.’ CRAN.

Sarup, P., Jensen, J., Ortersen, T., Henryon, M., & Sorensen, P. (2016). Increased prediction accuracy using a genomic feature model including prior information on quantitative trait locus regions in purebred Danish Duroc pigs. BMC Genomics, 17, 1–16

Schauer, N., Semel, Y., Balbo, I., Steinfath, M., Repsilber, D., Selbig, J., Pleban, T., Zamir, D., & Fernie, A.R. (2008). Mode of Inheritance of Primary Metabolic Traits in Tomato. The Plant Cell, 20, 509–523. https://doi.org/10.1105/tpc.107.056523

Soltis, N.E., & Kliebenstein, D.J. (2015). Natural variation of plant metabolism: Genetic mechanisms, interpretive caveats, and evolutionary and mechanistic insights. Plant Physiology, 169, 1456–1468. https://doi.org/10.1104/pp.15.01108

Stewart, D., & McDougall, G. (2014). Oat agriculture, cultivation and breeding targets: Implications for human nutrition and health. British Journal of Nutrition, 112, S50–S57. https://doi.org/10.1017/S0007114514002736

Tsugawa, H., Kind, T., Nakabayashi, R., Yukihira, D., Tanaka, W., Cajka, T., Saito, K., Fiehn, O., & Arita, M. (2016). Hydrogen Rearrangement Rules: Computational MS/MS Fragmentation and Structure Elucidation Using MS-FINDER Software. Analytical Chemistry, 88, 7946–7958. https://doi.org/10.1021/acs.analchem.6b00770

Turner-Hissong, S.D., Bird, K.A., Lipka, A.E., King, E.G., Beissinger, T.M., & Angelovici, R. (2020). Genomic Prediction Informed by Biological Processes Expands Our Understanding of the Genetic Architecture Underlying Free Amino Acid Traits in Dry *Arabidopsis* Seeds. G3 Genes|Genomes|Genetics, 10, 4227–4239. https://doi.org/10.1534/g3.120.401240

Wager, A., & Li, X. (2018). Exploiting natural variation for accelerating discoveries in plant specialized metabolism. Phytochemistry Reviews, 17, 17–36. https://doi.org/10.1007/s11101-017-9524-2

Wang, H., Cimen, E., Singh, N., & Buckler, E. (2020). Deep learning for plant genomics and crop improvement. Current Opinion in Plant Biology, 54, 34–41. https://doi.org/10.1016/j.pbi.2019.12.010

Zhou, S., Kremling, K.A., Bandillo, N., Richter, A., Zhang, Y.K., Ahern, K.R., Artyukhin, A.B., Hui, J.X., Younkin, G.C., Schroeder, F.C., Buckler, E.S., & Jander, G. (2019). Metabolome-scale genome-wide association studies reveal chemical diversity and genetic control of maize specialized metabolites. Plant Cell, 31, 937–955. https://doi.org/10.1105/tpc.18.00772

Zhu, G., Gou, J., Klee, H., & Huang, S. (2019). Next-Gen Approaches to Flavor-Related Metabolism. Annual Review of Plant Biology, 70, 187–212

Zhu, G., Wang, S., Huang, Z., Zhang, S., Liao, Q., Zhang, C., Lin, T., Qin, M., Peng, M., Yang, C., Cao, X., Han, X., Wang, X., van der Knaap, E., Zhang, Z., Cui, X., Klee, H., Fernie, A.R., Luo, J., & Huang, S. (2018). Rewiring of the Fruit Metabolome in Tomato Breeding. Cell, 172, 249–261.e12. https://doi.org/10.1016/j.cell.2017.12.019

