## Supporting Information for "Generalizable approaches for genomic prediction of metabolites in plants"

### SUPPORTING INFORMATION CONTENTS

**File S1**. Validation germplasm panel metabolite information

**Figure S1**. Linear regressions of metabolite heritability by retention time and molecular mass.

**Figure S2**. Number of metabolites with significant GWAS results by instrument type.

**Figure S3**. The number of metabolites and SNPs contributing to kernels.

**Figure S4**. Correlation between off-diagonal elements in metabolite kernels.

**Figure S5**. Mean Euclidean distance between metabolites that contribute to metabolite kernels.

**Figure S6**. Genomic distribution of metabolite GWAS results and the general kernels.

**Figure S7**. Genomic distribution of metabolite GWAS results and the lipid kernels.

**Figure S8**. Validation panel metabolite genomic heritability and relationship to retention time.

**Figure S9**. Mean cross-fold validation accuracy of metabolites compared to genomic heritability

**Figure S10**. Metabolites where any two-kernel metabolite model improved or reduced genomic prediction accuracy over GBLUP.

**Figure S11**. Percent genetic variation attributed to the metabolite kernel compared to metabolite genomic heritability.

**Table S1**. Mean metabolite heritability by ClassyFire superclass groups for the discovery panel.

**Table S2**. Number of metabolites in each metabolite kernel.

**Table S3**. Number of SNPs in each metabolite kernel.

**Table S4**. Spearman's rank correlation of metabolite heritability across validation panel environments.

**Table S5**. Validation panel LC-MS metabolite ClassyFire 'Superclass' count and heritability.

**Table S6.** Validation panel linear regression between retention time and heritability

**Table S7**. Comparison of percent genetic variance (percent heritability) by metabolite kernels to mGWAS random kernels by instrument, model and environment.

**Table S8**. Significant differences in prediction accuracy compared to GBLUP for seven specialized metabolites.

**Table S9**. Coefficient of variation in retention time of LC-MS lipids by Class.

**File S1.** Deregressed BLUPs of the metabolites in the validation germplasm panel, and descriptions of metabolites are provided in the file: Validation_Panel_Metabolite_Data..xlsx

**Figure S1**. Linear regressions of metabolite heritability by retention time and molecular mass for (a,c) all metabolites and (b,d) with the lowest heritability metabolites removed ($h^2$>0.01) for the discovery panel. The blue and red points, lines and labels indicate the LC and GC metabolites, respectively. The results of ANOVA analysis for retention time and molecular mass for all metabolites are as follows: retention time LC-MS: $F_{1,1065}$ = 292.5, $p$ < 2.2e-16; retention time GC-MS: $F_{1,599}$ = 30.6, $p$ < 4.6e-08; molecular masses LC-MS: $F_{1,1065}$ = 9.0, $p$ = 0.003. The results of ANOVA analysis for retention time and molecular mass with metabolites with $h^2$>0.01 only are as follows: retention time, LC-MS: $F_{1,976}$ = 267.5, $p$ < 2.2e-16; retention time, GC-MS: $F_{1,431}$ = 18.9, $p$ < 1.74e-05; molecular mass, LC-MS: $F_{1,976}$ = 9.0, $p$ = 0.012.

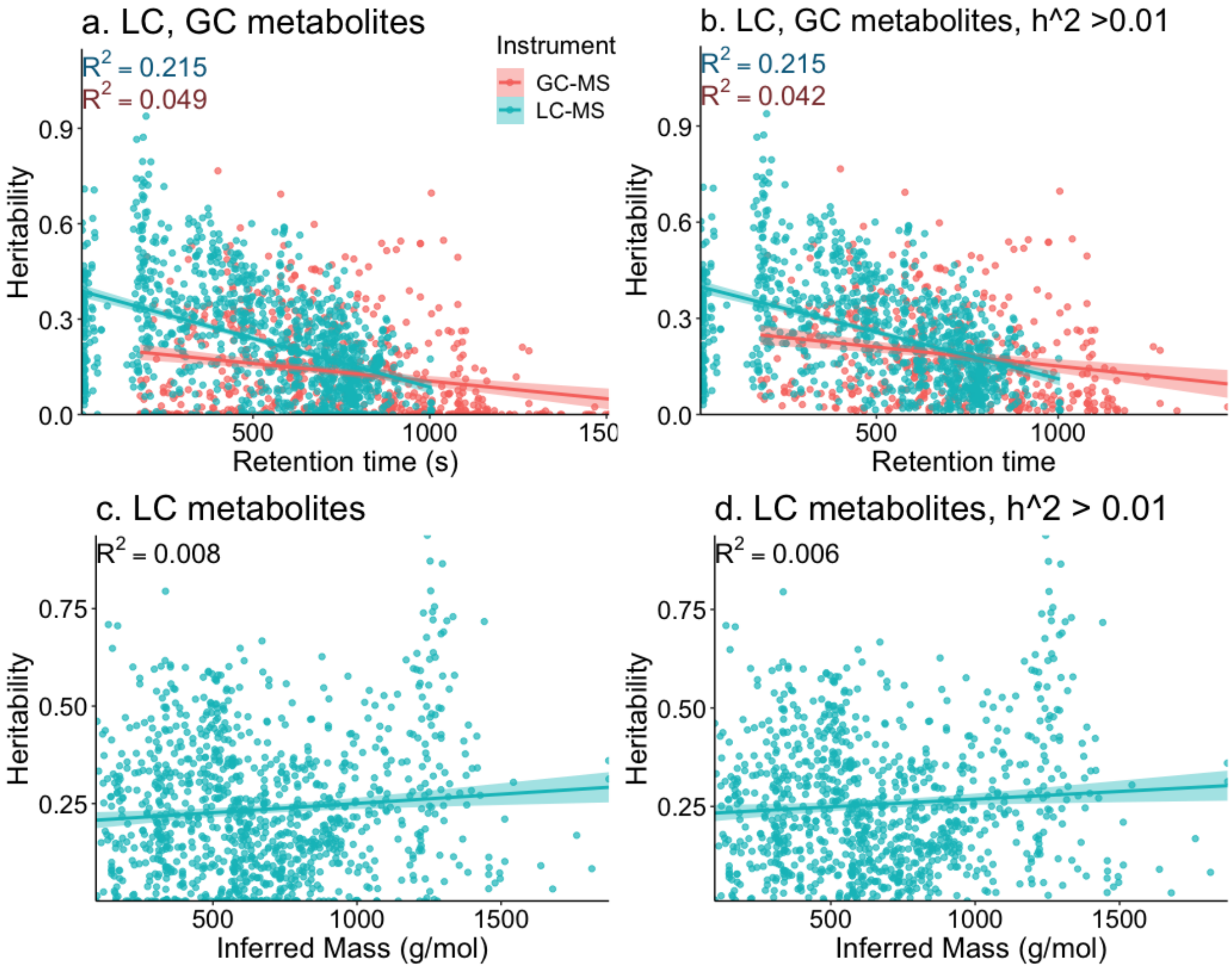

**Figure S2**. Number of metabolites with significant GWAS results by instrument type (LC-MS, GC-MS).

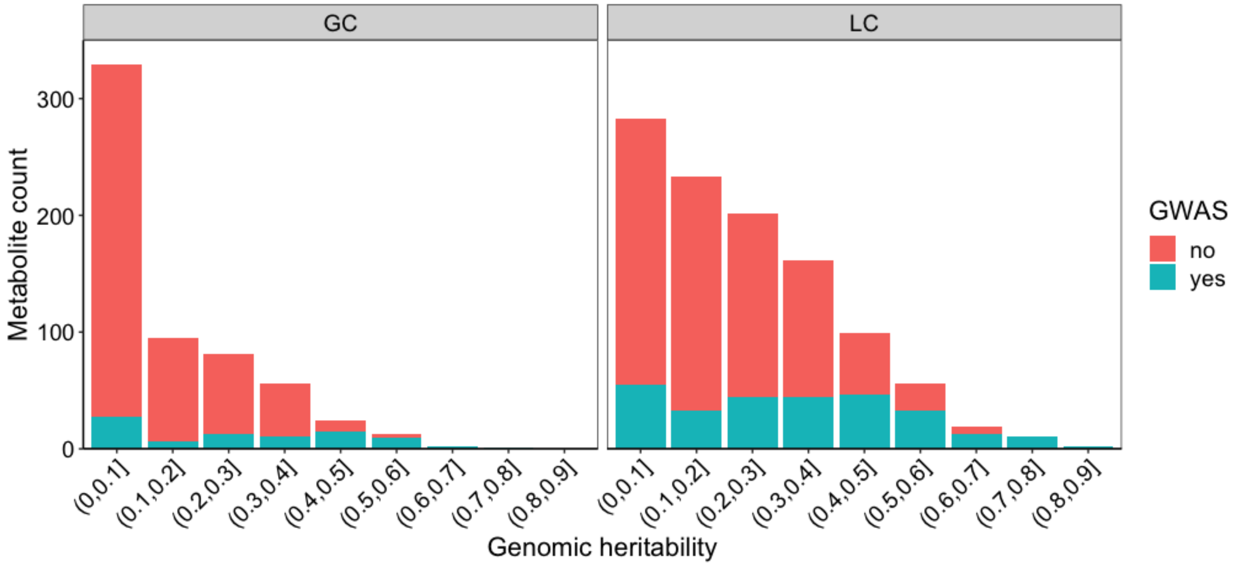

**Figure S3** The number of (a, c) metabolites and (b, d) SNPs contributing to the (a,b) general kernels 'Any3', 'LCGC2', 'LC4' and 'GC2' and (c,d) lipid specific kernels of 'Lipid', 'MVA' and 'MEP'. The Lipid kernel shared 65 metabolites and 378 SNPs with the LC4 kernel (13 metabolites and 272 SNPs were not shared)

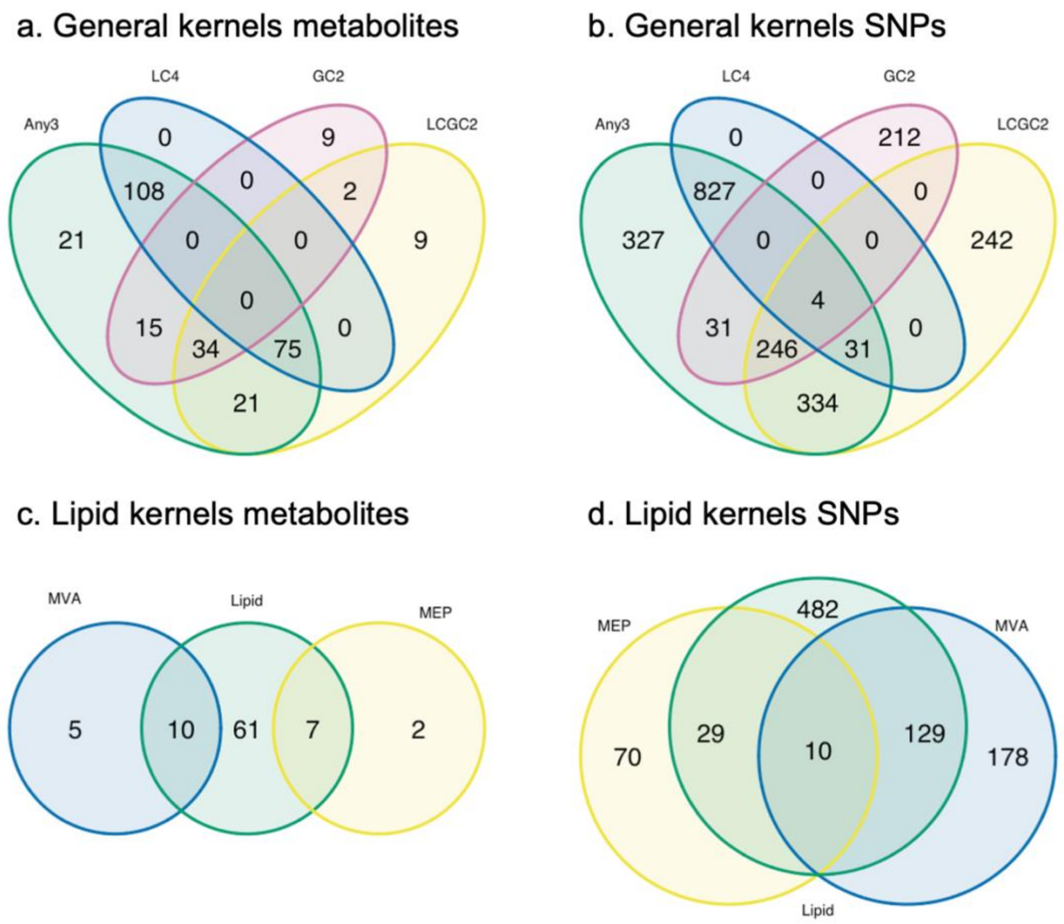

**Figure S4**. Correlation between off-diagonal elements in metabolite kernels.

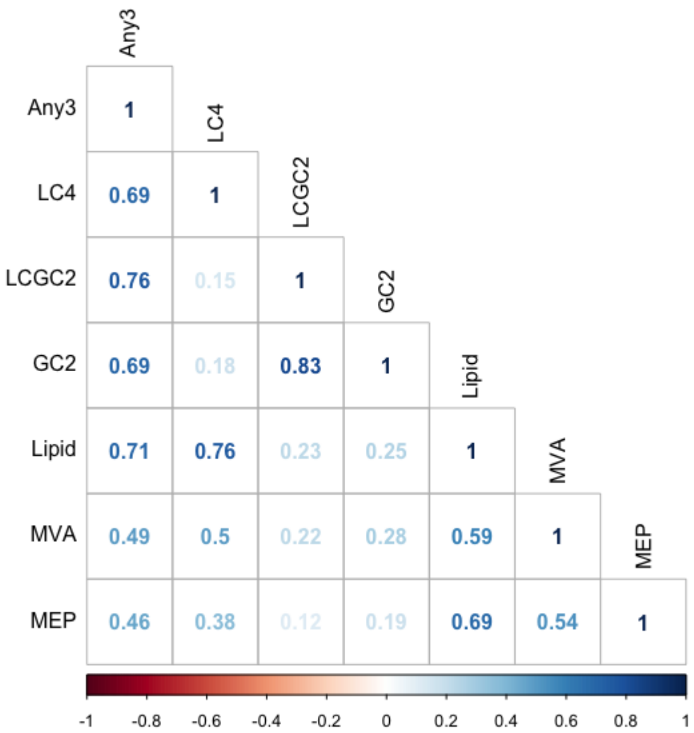

**Figure S5**. Mean Euclidean distance between metabolites that contribute to metabolite kernels developed in the discovery panel. Distance measures of kernels were compared to all metabolites (“All”) by Wilcoxon signed rank test. The * indicates a *p*-value less than the Bonferroni cutoff per plot, and ** and *** indicate p < 1e-4, and p<1e-6, respectively.

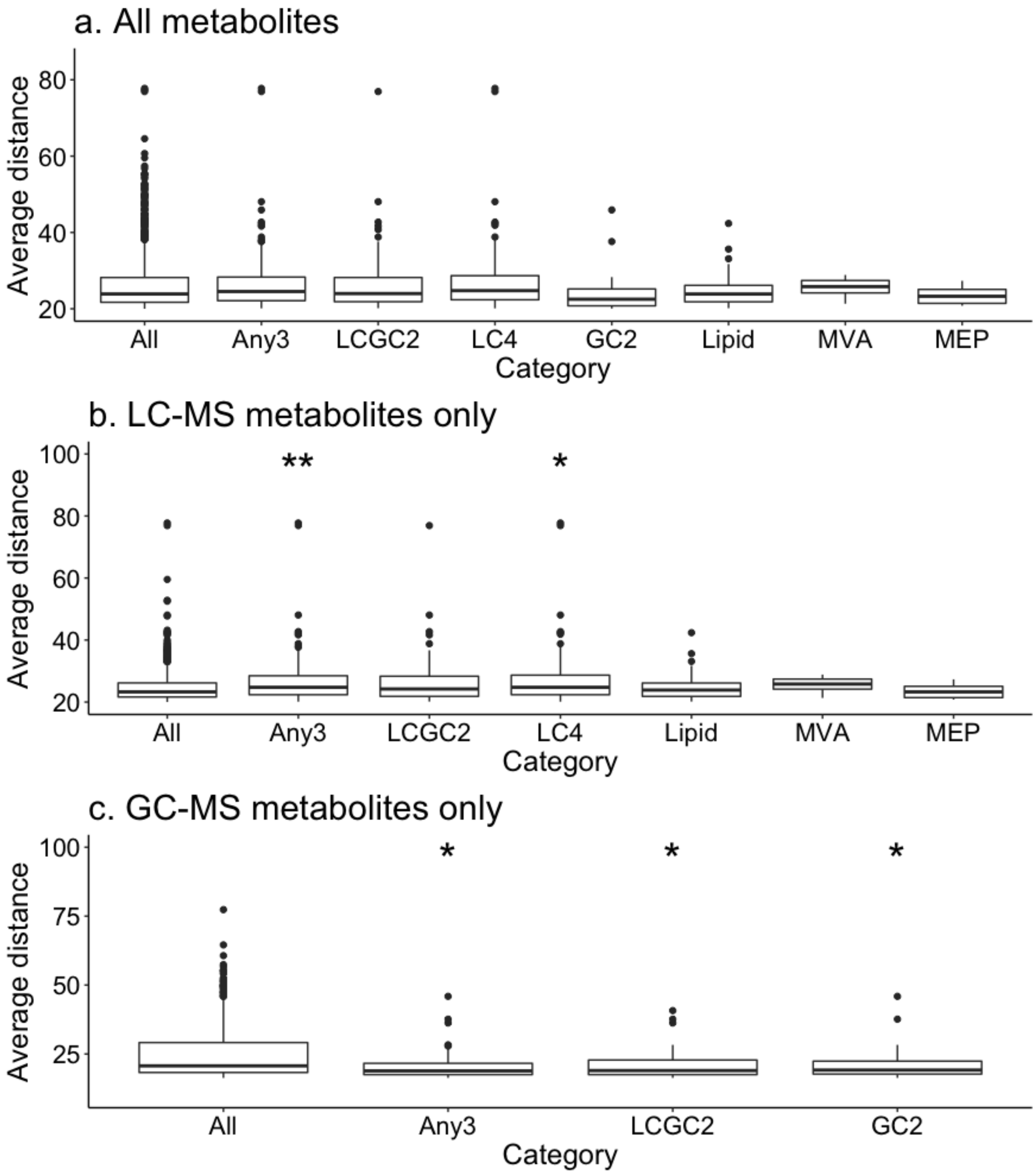

**Figure S6.** Genomic distribution of metabolite GWAS results and the general kernels (a) all GWAS results, (b) 'Any3' kernel, (c) 'LCGC2' kernel, (d) 'LC4' kernel, and (e) 'GC2' kernel. The count of GWAS results are summed and plotted by 10 Mb bins.

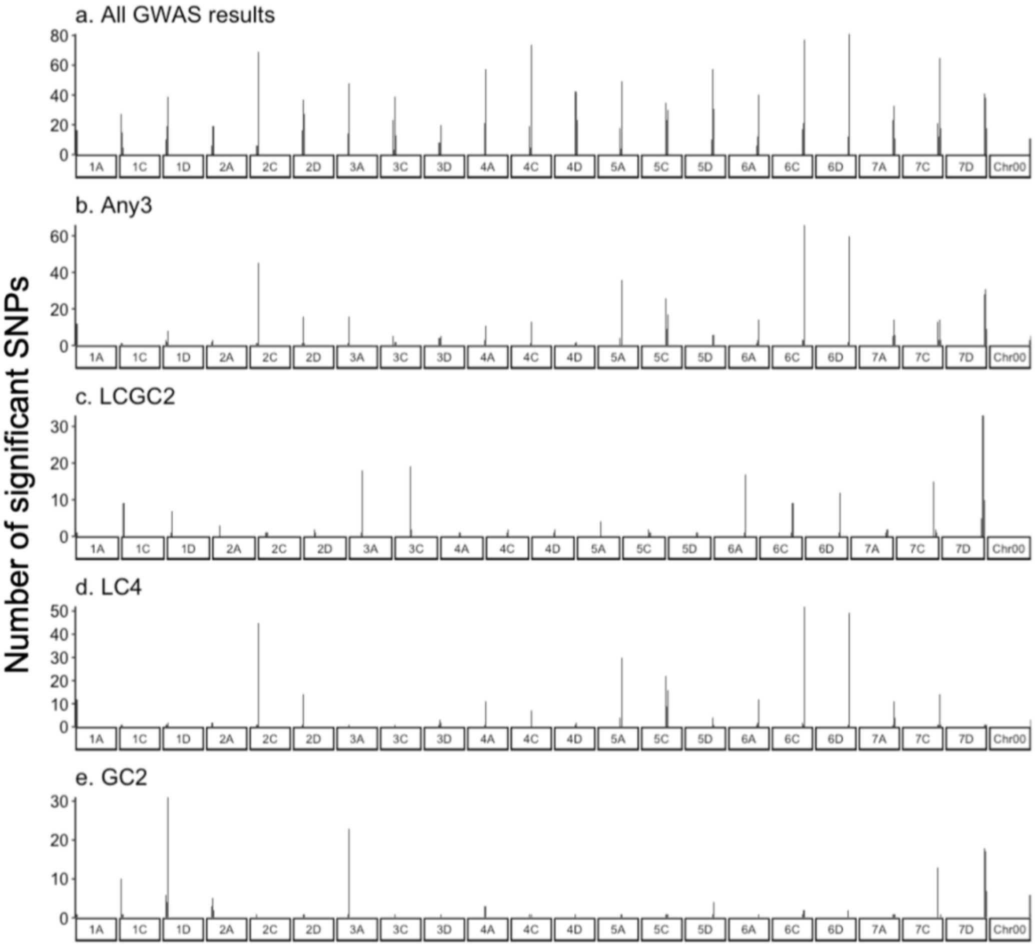

**Figure S7.** Genomic distribution of metabolite GWAS results and the lipid kernels (a) all lipids, 'Lipid', and the two terpenoid pathway kernels (b) MEP kernel, and (c) MVA kernel. The count of GWAS results are summed and plotted by 10 Mb bins.

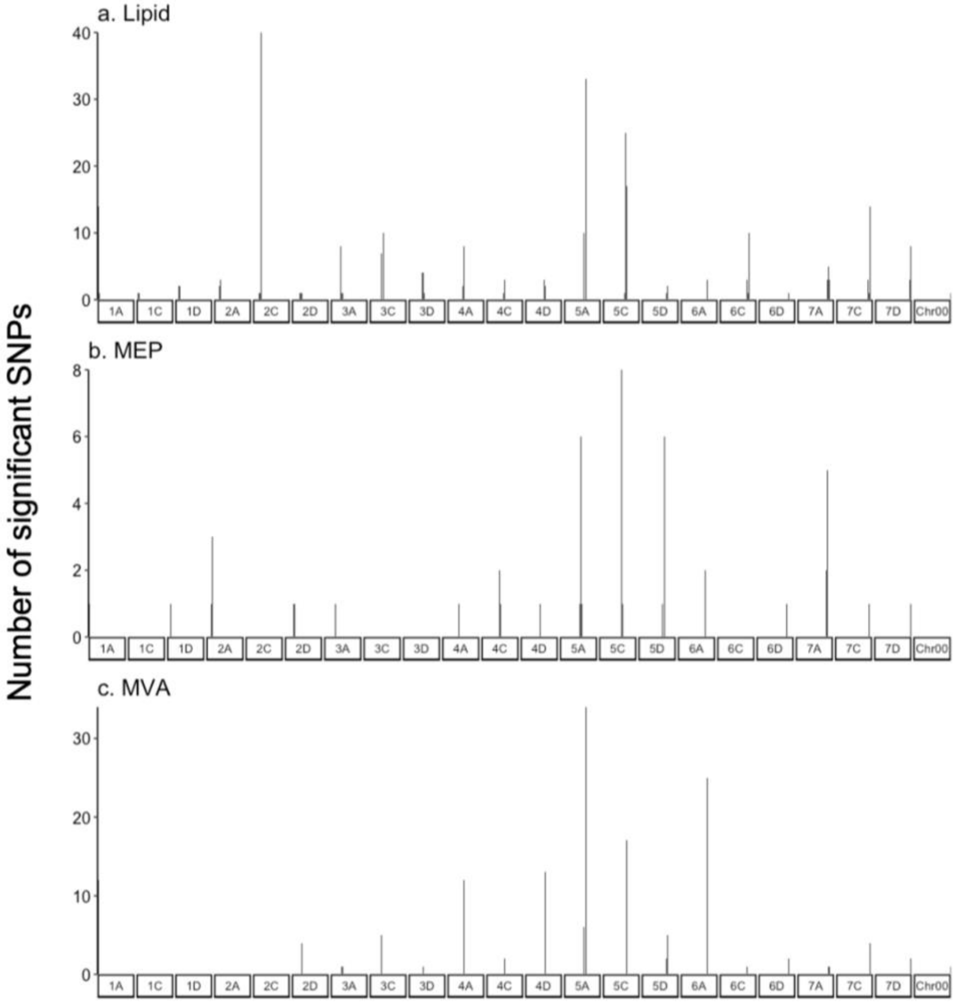

**Figure S8**. Validation panel metabolite genomic heritability distribution and (b, d, f) relationship between heritability and retention time by environment. For all plots, the instrument class (LC-MS, or GC-MS) is denoted by color (blue, red, respectively). For genomic heritability for (a) Minnesota ("MN"), (c) South Dakota ("SD") and (e) Wisconsin ("WI"), the solid line indicates the mean and dashed line indicates the median genomic heritability by instrument class. The linear regression of metabolite heritability by retention time for (b) MN, (d) SD and (f) WI is given along with the model coefficient of determination ($R^2$). The regression results are given in Table S4.

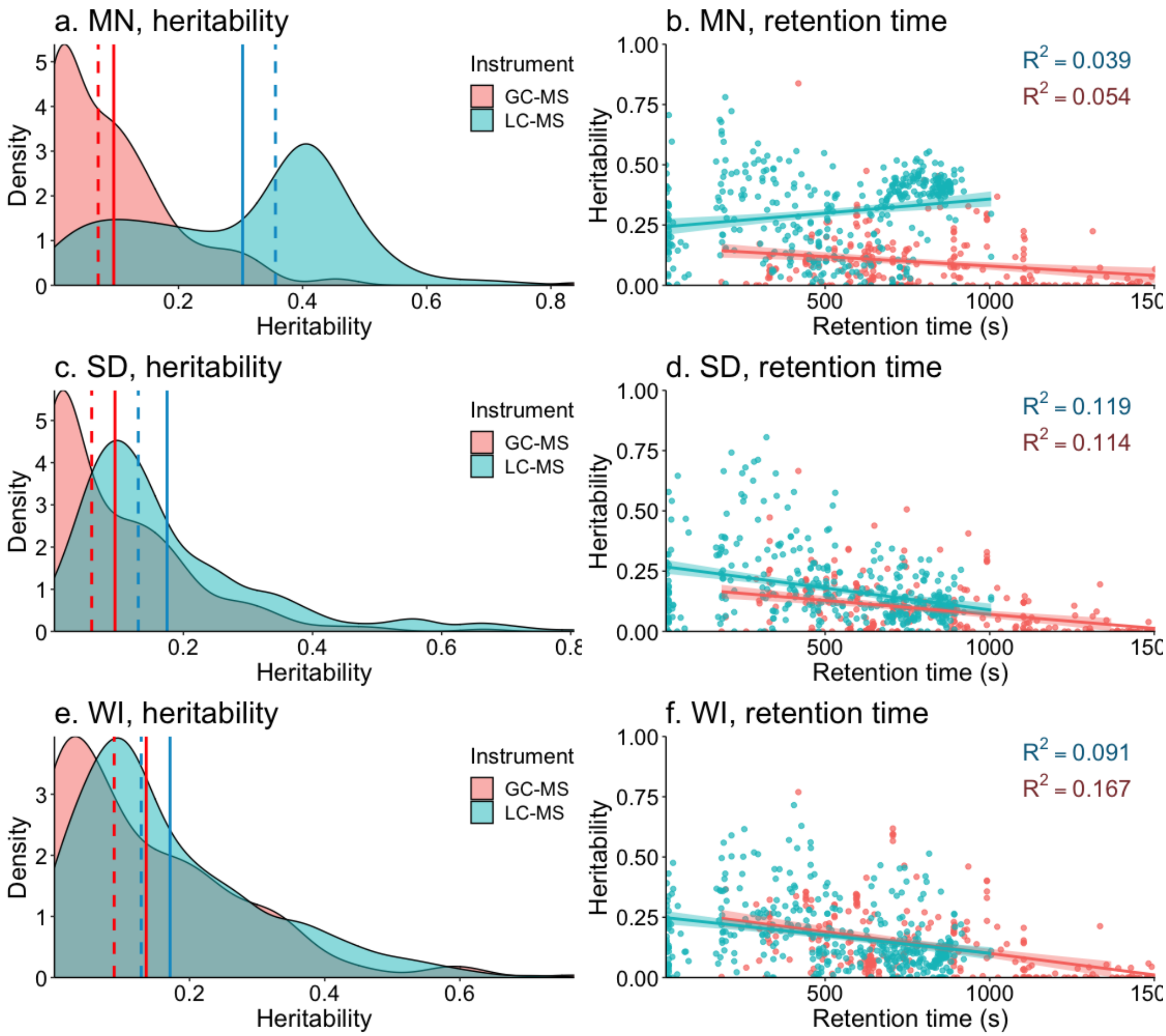

**Figure S9**. Mean cross-fold validation accuracy (r) of metabolites as compared to genomic heritability for (a.) LC-MS (*n*=396) and (b.) GC-MS (*n*=243) metabolites by environment (Minnesota, “MN”; South Dakota, “SD” and Wisconsin, “WI”) for all metabolite models combined. The coefficient of determination is reported in each individual plot.

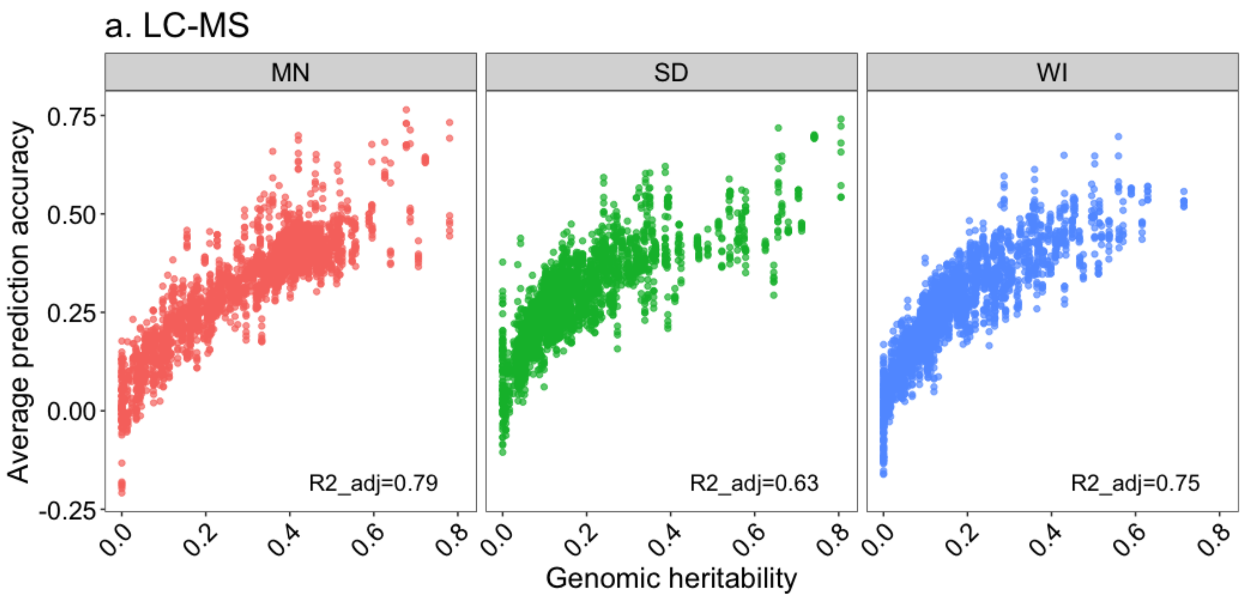


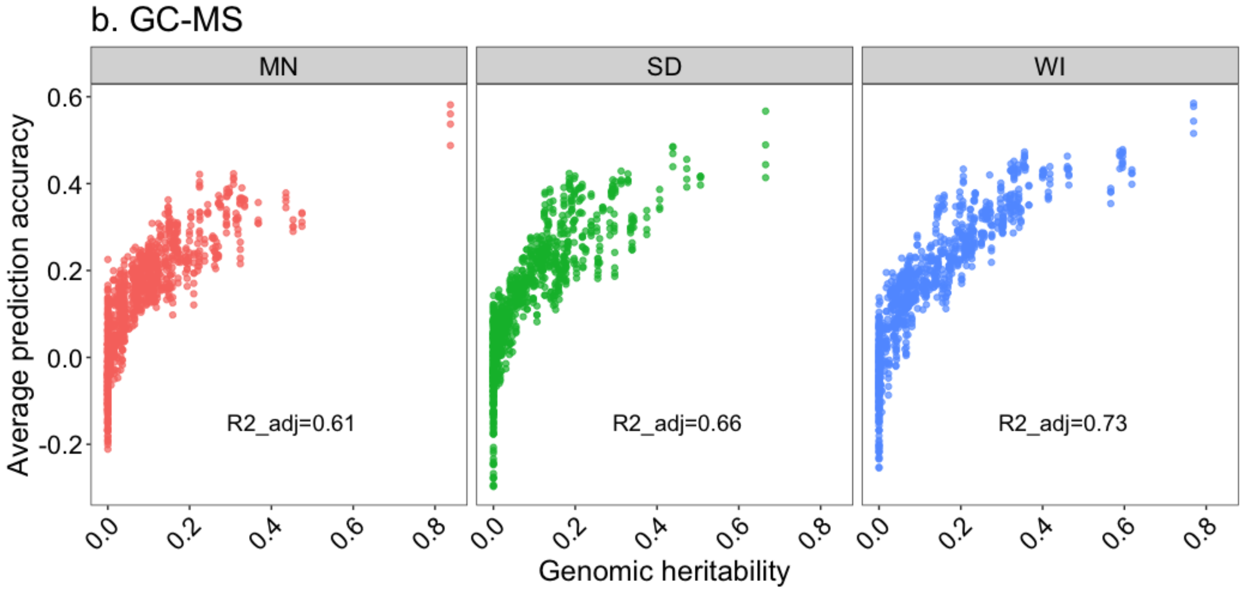

**Figure S10.** Number of metabolites where any two-kernel metabolite model improved or reduced genomic prediction accuracy over GBLUP and the degree to which the metabolites were shared by environment (Minnesota, "MN"; South Dakota, "SD" and Wisconsin, "WI").

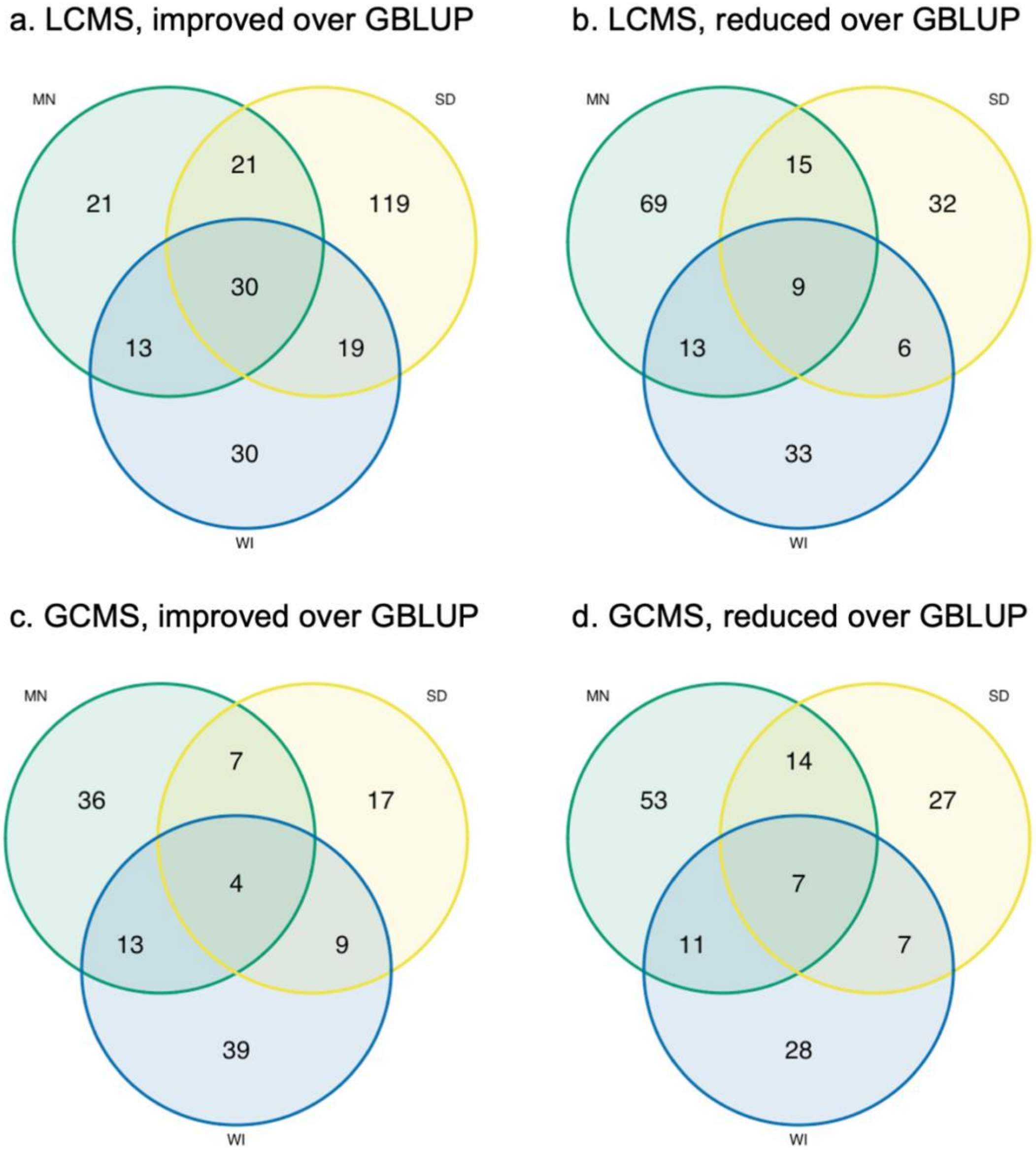

**Figure S11**. Percent genetic variation attributed to the metabolite kernel as compared to metabolite genomic heritability for (a.) LC-MS (*n*=397) and (b.) GC-MS (*n*=243) metabolites by environment (Minnesota, "MN"; South Dakota, "SD" and Wisconsin, "WI"). The coefficient of determination is reported in each individual plot.

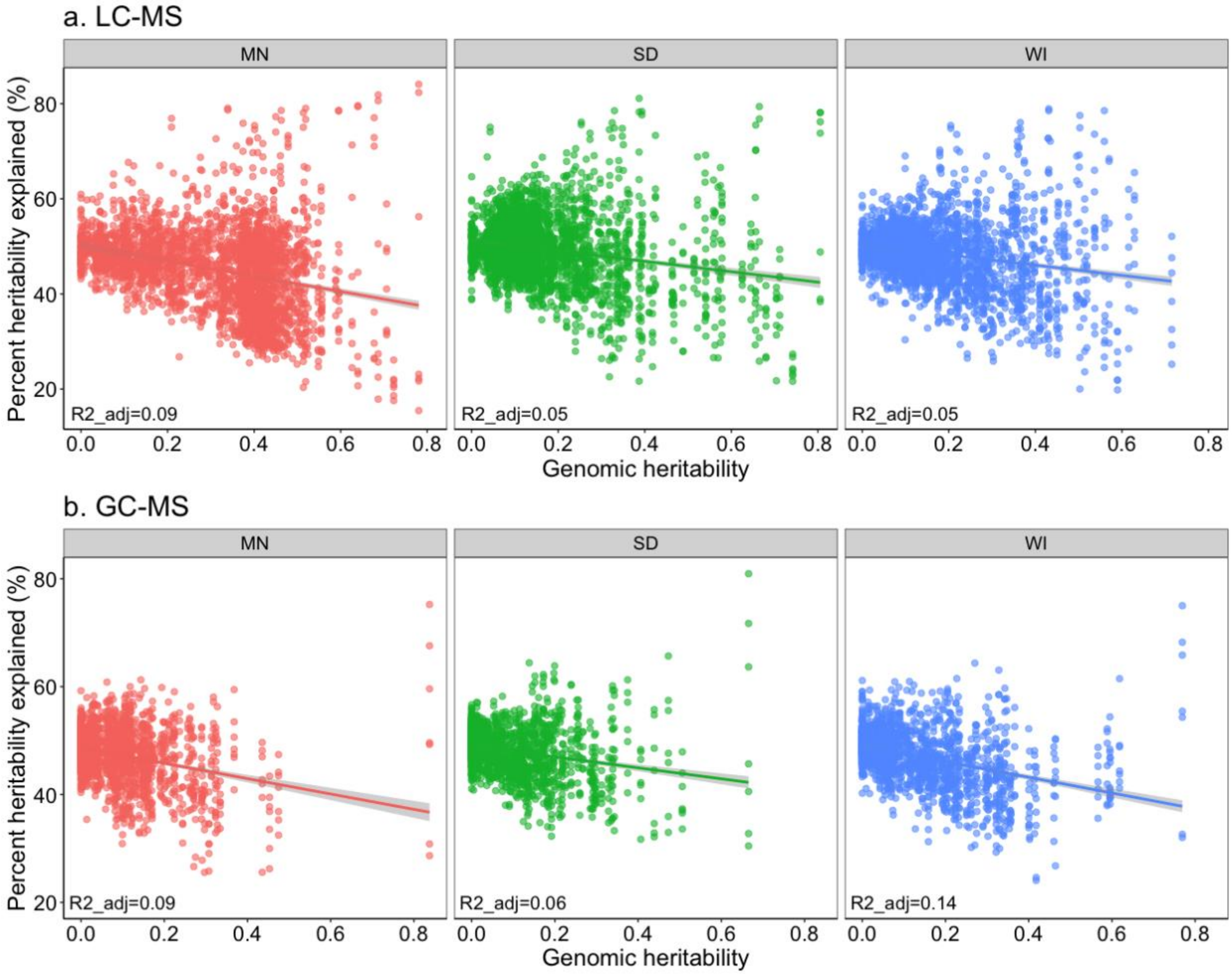

**Table S1**. Mean metabolite heritability by ClassyFire superclass groups for the discovery panel.

| Classification | LC | GC |
|---|---|---|
| Overall | 0.23 | 0.13 |
| Lipids and lipid-like molecules | 0.21 | 0.15 |
| Organoheterocyclic compounds | 0.26 | 0.29 |
| Phenylpropanoids and polyketides | 0.26 | 0.03 |
| Organic acids and derivatives | 0.29 | 0.35 |
| Organic oxygen compounds | 0.23 | 0.27 |
| Not classified | 0.26 | 0.12 |

**Table S2**. Number of metabolites in each kernel, where total metabolites ("Total") and metabolites with a GWAS result ("GWAS") are presented for comparison.

| Kernel | Number LC | Number GC | Total Number |
|---|---|---|---|
| Total | 1067 | 601 | 1668 |
| GWAS | 282 | 86 | 368 |
| Any3 | 220 | 54 | 274 |
| LCGC2 | 95 | 46 | 141 |
| LC4 | 183 | 0 | 183 |
| GC2 | 0 | 60 | 60 |
| Lipid | 78 | 0 | 78 |
| MEP | 9 | 0 | 9 |
| MVA | 15 | 0 | 15 |

**Table S3**. Number of SNPs in each shared GWAS kernel before and after adding SNPs that were in LD ($r^2$>0.5).

| Kernel | Number of SNPs | Number of SNPs with those in LD |
|---|---|---|
| Any3 | 1800 | 5311 |
| LC _ GC _2 | 857 | 2593 |
| LC _ 4 | 862 | 2596 |
| GC2 | 493 | 2075 |
| Lipid | 650 | 2440 |
| MEP | 109 | 620 |
| MVA | 317 | 999 |

**Table S4**. Spearman's rank correlation of metabolite heritability across environments (Minnesota, "MN", South Dakota, "SD", and Wisconsin, "WI") in the validation panel.

| Instrument | Environment | SD | WI |
|---|---|---|---|
| LC | MN | rho=0.20, *p*=8.4e-05 | rho=0.16, *p*=1.3e-05=3 |
| LC | SD | | rho=0.61, *p*<2.2e-16 |
| GC | MN | rho=0.55, *p*<2.2e-16 | rho=0.59, *p*<2.2e-16 |
| GC | SD | | rho=0.69, *p*<2.2e-16 |

**Table S5**. Validation panel LC-MS metabolite ClassyFire 'Superclass' count and heritability by category and environment (Minnesota, "MN"; South Dakota, "SD" and Wisconsin, "WI").

| Classification | Count | MN $h^2$ | SD $h^2$ | WI $h^2$ |
|---|---|---|---|---|
| Lipids and lipid-like molecules | 91 | 0.27 | 0.16 | 0.17 |
| Organoheterocyclic compounds | 9 | 0.25 | 0.22 | 0.20 |
| Phenylpropanoids and polyketides | 6 | 0.30 | 0.34 | 0.27 |
| Organic acids and derivatives | 9 | 0.16 | 0.20 | 0.18 |
| Organic oxygen compounds | 3 | 0.49 | 0.21 | 0.20 |
| Other | 16 | 0.25 | 0.19 | 0.17 |
| Not classified | 263 | 0.32 | 0.17 | 0.17 |

**Table S6**. Validation panel results from linear regression between retention time ("RT") and heritability

| Instrument | Environment | RT coefficient sign | Model results |
|---|---|---|---|
| LC-MS | MN | positive | F(1,394)=16.04, **p=7.41e-05** |
| LC-MS | SD | negative | F(1,395)=53.45, **p=1.49e-12** |
| LC-MS | WI | negative | F(1,395)=39.61, **p=8.27e-10** |
| GC-MS | MN | negative | F(1,241)=13.71, **p=2.64e-04** |
| GC-MS | SD | negative | F(1,241)=31.12, **p=2.64e-04** |
| GC-MS | WI | negative | F(1,241)=48.29, **p=3.42e-11** |

**Table S7.** Comparison of percent genetic variance (percent heritability) by metabolite kernels to mGWAS random kernels by instrument, model and environment. Significance tests are paired Wilcox signed rank tests, where significance identifiers are: * p<0.05, **p<0.001; *** p<1e-6

| Instrument | Model | Env | Metabolite kernel | | Random mGWAS kernel | | Wilcoxon Rank | Direction |
|---|---|---|---|---|---|---|---|---|
| | | | *n* SNPs | mean h2 | *n* SNPs | mean h2 | | |
| LC-MS | Any3 | MN | 5311 | 46.9 | 6660 | 46.6 | ns | better |
| | | SD | 5311 | 51.1 | 6660 | 48.9 | *** | better |
| | | WI | 5311 | 50.0 | 6660 | 48.9 | * | better |
| | LCGC2 | MN | 2593 | 39.8 | 2733 | 46.8 | *** | worse |
| | | SD | 2593 | 50.1 | 2733 | 48.9 | *** | better |
| | | WI | 2593 | 47.9 | 2733 | 48.8 | ns | worse |
| | LC4 | MN | 2596 | 49.5 | 2733 | 46.8 | *** | better |
| | | SD | 2596 | 49.7 | 2733 | 48.9 | * | better |
| | | WI | 2596 | 49.5 | 2733 | 48.8 | ns | better |
| | GC2 | MN | 2075 | 41.0 | 2733 | 46.8 | *** | worse |
| | | SD | 2075 | 45.3 | 2733 | 48.9 | *** | worse |
| | | WI | 2075 | 45.1 | 2733 | 48.8 | *** | worse |
| | Lipid | MN | 2440 | 48.7 | 2733 | 46.8 | *** | better |
| | | SD | 2440 | 50.1 | 2733 | 48.9 | ** | better |
| | | WI | 2440 | 49.3 | 2733 | 48.8 | ns | better |
| | MEP | MN | 620 | 41.7 | 697 | 46.7 | *** | worse |
| | | SD | 620 | 47.6 | 697 | 49.0 | * | worse |
| | | WI | 620 | 49.7 | 697 | 48.7 | * | better |
| | MVA | MN | 999 | 49.5 | 1336 | 46.7 | *** | better |
| | | SD | 999 | 51.5 | 1336 | 49.0 | *** | better |
| | | WI | 999 | 48.0 | 1336 | 48.7 | * | worse |
| GC-MS | Any3 | MN | 5311 | 48.5 | 6660 | 49.7 | ** | worse |
| | | SD | 5311 | 49.0 | 6660 | 49.8 | * | worse |
| | | WI | 5311 | 48.0 | 6660 | 49.5 | ** | worse |
| | LCGC2 | MN | 2593 | 46.1 | 2733 | 49.6 | *** | worse |
| | | SD | 2593 | 49.6 | 2733 | 49.8 | ns | worse |
| | | WI | 2593 | 47.2 | 2733 | 49.4 | *** | worse |
| | LC4 | MN | 2596 | 48.2 | 2733 | 49.6 | * | worse |
| | | SD | 2596 | 47.8 | 2733 | 49.8 | *** | worse |
| | | WI | 2596 | 47.5 | 2733 | 49.4 | *** | worse |
| | GC2 | MN | 2075 | 47.1 | 2733 | 49.6 | *** | worse |
| | | SD | 2075 | 48.5 | 2733 | 49.8 | ** | worse |
| | | WI | 2075 | 47.9 | 2733 | 49.4 | ** | worse |
| | Lipid | MN | 2440 | 49.4 | 2733 | 49.6 | ns | worse |
| | | SD | 2440 | 48.3 | 2733 | 49.8 | *** | worse |
| | | WI | 2440 | 47.8 | 2733 | 49.4 | ** | worse |
| | MEP | MN | 620 | 46.1 | 697 | 49.6 | *** | worse |
| | | SD | 620 | 46.1 | 697 | 49.7 | *** | worse |
| | | WI | 620 | 46.0 | 697 | 49.4 | *** | worse |
| | MVA | MN | 999 | 45.6 | 1336 | 49.7 | *** | worse |
| | | SD | 999 | 47.1 | 1336 | 49.7 | *** | worse |
| | | WI | 999 | 44.9 | 1336 | 49.5 | *** | worse |

**Table S8.** Instances in which prediction accuracy of two kernel model was significantly greater than GBLUP for seven specialized metabolites ($p_{BONF}$<0.05 metabolome-wide). The p-value is from paired Wilcoxon tests and unadjusted. Genomic heritability estimates are given by environment ("env"). The metabolite abbreviations are as follows: AVN_A, avenanthramide A; AVN_B, avenanthramide B; AEC_A1, AEC_A2, avenacin A1; AOS_A, avenacoside A; AOS_dA, 26-Desglucoavenacoside A; AOS_B, avenacoside B

| Type | Metabolite | model | p | Env | Env_h2 |
|---|---|---|---|---|---|
| Avenanthramides | AVN_A | MVA | 7.3E-07 | SD | 0.14 |
| | AVN_A | MEP | 5.2E-09 | SD | 0.14 |
| | AVN_B | MEP | 6.8E-08 | SD | 0.14 |
| Avenacins | AEC_A1 | Any3 | 5.8E-09 | MN | 0.46 |
| | AEC_A2 | Any3 | 1.1E-06 | MN | 0.33 |
| | AEC_A1 | LC4 | 1.3E-08 | MN | 0.46 |
| | AEC_A2 | LC4 | 6.8E-08 | MN | 0.33 |
| | AEC_A1 | Lipid | 2.5E-09 | MN | 0.46 |
| | AEC_A2 | Lipid | 1.8E-07 | MN | 0.33 |
| | AEC_A2 | Lipid | 1.3E-08 | WI | 0.37 |
| | AEC_A1 | MVA | 3.7E-09 | MN | 0.46 |
| | AEC_A2 | MVA | 2.5E-08 | MN | 0.33 |
| | AEC_A2 | MVA | 6.9E-09 | WI | 0.37 |
| Avenacosides | AOS_B | LCGC2 | 6.4E-08 | WI | 1.00E-09 |
| | AOS_dA | Lipid | 1.3E-09 | MN | 0.26 |
| | AOS_dA | Lipid | 5.7E-07 | SD | 0.08 |
| | AOS_dA | MVA | 4.1E-06 | SD | 0.08 |
| | AOS_B | MVA | 5.8E-09 | SD | 0.01 |
| | AOS_B | MVA | 3.9E-10 | WI | 1.00E-09 |
| | AOS_A | MEP | 3.9E-06 | SD | 0.16 |
| | AOS_A | MEP | 2.0E-06 | WI | 0.08 |
| | AOS_dA | MEP | 1.6E-07 | MN | 0.26 |
| | AOS_dA | MEP | 1.7E-07 | SD | 0.08 |
| | AOS_B | MEP | 3.5E-09 | WI | 1.00E-09 |

**Table S9.** Coefficient of variation ("CV") in retention time (s) of LC-MS lipids by Class. The number of metabolites in each Class is given by 'n'.

| Class | Color code | n | CV |
|---|---|---|---|
| Glycerolipids ("GL") | Black | 36 | 20.6 |
| Fatty Acyls ("FA") | Green | 17 | 22.6 |
| Prenol lipids ("PL) | Red | 14 | 35.3 |
| Steroids and steroid derivatives ("ST") | Purple | 12 | 39.2 |
| Glycerophospholipids ("GP") | Blue | 9 | 70.7 |
| Sphingolipids ("SP") | Grey | 2 | 5.9 |
| Saccharolipids ("SA") | Grey | 1 | NA |
